# The *Pseudomonas aeruginosa* LasR quorum-sensing receptor balances ligand selectivity and sensitivity

**DOI:** 10.1101/269001

**Authors:** Amelia R. McCready, Jon E. Paczkowski, Brad R. Henke, Bonnie L. Bassler

**Affiliations:** The Department of Molecular Biology, Princeton University, Princeton, NJ 08544; Opti-Mol Consulting, LLC, Cary, NC 27513; Howard Hughes Medical Institute, Chevy Chase, MD 20815

**Author notes:** To whom correspondence should be addressed: Prof. Bonnie L. Bassler, Department of Molecular Biology, Princeton University, 329 Lewis Thomas Laboratory, Princeton, NJ 08544.

**Keywords:** crystal structure, LasR, *Pseudomonas aeruginosa*, quorum sensing, receptor specificity

## Abstract

Quorum sensing is a cell-cell communication process that bacteria use to orchestrate group behaviors. Quorum sensing is mediated by extracellular signal molecules called autoinducers. Autoinducers are often structurally similar, raising questions concerning how bacteria distinguish among them. Here, we use the *Pseudomonas aeruginosa* LasR quorum-sensing receptor to explore receptor sensitivity and selectivity. Alteration of LasR amino acid S129 increases ligand selectivity and decreases ligand sensitivity. Conversely, the L130F mutation enhances LasR sensitivity while reducing selectivity. We solve crystal structures of LasR ligand binding domains complexed with non-cognate autoinducers. Comparison to existing structures reveals that ligand selectivity/sensitivity is mediated by a flexible loop adjacent to the ligand binding site. We show that *P. aeruginosa* harboring LasR variants with modified selectivity or sensitivity exhibit altered quorum-sensing responses. We suggest that an evolutionary trade-off between ligand selectivity and sensitivity enables LasR to optimally regulate quorum-sensing traits.

## Introduction

Quorum sensing is a cell-cell communication process that enables bacteria to collectively control behavior (reviewed in Papenfort & Bassler, 2016). Quorum sensing relies on the production, release, and detection of extracellular signal molecules called autoinducers (Albus, Pesci, RunyenJanecky, West, & Iglewski, 1997; Engebrecht, Nealson, & Silverman, 1983; Latifi et al., 1995). At low cell density, autoinducer concentration is low, and bacteria act as individuals. As cell density increases, autoinducer concentration also rises. Under this condition, autoinducers bind to cognate receptors, initiating the population-wide regulation of genes underlying collective behaviors.

Many species of Gram-negative bacteria use acylated homoserine lactones [HSLs] as autoinducers (Brint & Ohman, 1995; Cao & Meighen, 1989; Eberhard et al., 1981; Hanzelka et al., 1999; Pearson et al., 1994). HSL autoinducers possess identical lactone head groups, but they vary in acyl tail length and decoration. The tail modifications promote specificity between particular HSL autoinducers and partner receptors (Churchill & Chen, 2011). There are two kinds of HSL autoinducer receptors. First, there are LuxR-type receptors, which are cytoplasmic HSL-binding transcription factors that possess variable ligand binding domains [LBD] and well-conserved helix-turn-helix DNA binding domains [DBD] (Nasser & Reverchon, 2007; Vannini et al., 2002). There are also LuxN-type receptors, which are membrane-spanning two-component signaling proteins that bind HSL ligands in their periplasmic regions and transduce information regarding ligand occupancy internally by phosphorylation/dephosphorylation cascades (Bassler, Wright, Showalter, & Silverman, 1993; Freeman, Lilley, & Bassler, 2000; reviewed in Papenfort & Bassler, 2016).

Ligand sensitivity and selectivity has been examined in the founding member of the LuxN receptor family from *Vibrio harveyi* (Ke, Miller, & Bassler, 2015). LuxN is exquisitely selective for its cognate autoinducer 3OHC_4_HSL. Specific amino acids were identified in the predicted LuxN transmembrane spanning region that confer selectivity for tail length and for tail decoration. Longer HSLs competitively inhibit LuxN, suggesting that, in mixed-species consortia, *V. harveyi* monitors the vicinity for competing species, and in response to their presence, exploits LuxN antagonism to delay the launch of its quorum-sensing behaviors, thus avoiding loss of expensive public goods to non-kin.

Some analyses of ligand preference in LuxR-type receptors have been performed. TraR from *Agrobacterium tumefaciens* excludes non-native HSLs (Hawver, Jung, & Ng, 2016; Vannini et al., 2002; You et al., 2006; Zhu & Winans, 2001), LasR from *Pseudomonas aeruginosa* detects several long chain HSLs (Gerdt et al., 2017), and SdiA from *Escherichia coli* is highly promiscuous and avidly responds to HSLs with variable chain lengths (Michael, Smith, Swift, Heffron, & Ahmer, 2001; Nguyen et al., 2015; Sitnikov, Schineller, & Baldwin, 1996). Comparison of structures of the LasR and TraR LBDs suggests that increased hydrogen bonding to the ligand in TraR, compared to LasR, accounts for the selectivity difference (Gerdt et al., 2017). Here, we systematically explore the LasR response to 3OC_12_HSL and non-native HSLs with respect to selectivity and sensitivity. We use mutagenesis to establish the amino acid determinants that enable LasR to discriminate between HSLs. We identify LasR S129 as the amino acid residue that, when altered, improves LasR selectivity for HSLs by restricting the set of HSLs capable of activation. Mutations at LasR S129, however, reduce overall affinity for HSL ligands. In contrast, we find that LasR L130F exhibits diminished ligand selectivity, responding to a broader set of HSLs than wildtype LasR, with enhanced sensitivity. Altering LasR sensitivity or selectivity affects the timing and strength of quorum-sensing control of *P. aeruginosa* behaviors. Finally, we solve crystal structures of the LasR LBD L130F bound to non-native autoinducers to establish the structural basis underlying ligand selectivity and sensitivity. We find that a flexible loop located near the ligand binding pocket promotes ligand promiscuity in the wildtype protein. This loop exists in SdiA, which is promiscuous, but not in TraR which is highly specific for its cognate ligand. We propose that there is a trade-off between ligand selectivity and sensitivity in LasR and evolution has established a balance between ligand discrimination and ligand sensitivity. Because LasR requires higher concentrations of non-cognate autoinducers than its cognate autoinducer for activation, this tradeoff could allow LasR to robustly respond to its own signal molecule, even in the presence of other bacteria that are producing HSL autoinducers. Nonetheless, *P. aeruginosa* would be capable of reacting to the presence of these other bacteria when it is outnumbered.

## Results

### LasR responds to multiple HSL autoinducers

To investigate the preference LasR displays for different HSLs, we employed a plasmid reporter system in which transcription from a LasR-controlled promoter fused to luciferase (p*lasB-lux*) was assessed in *E. coli*. Arabinose-inducible *lasR* was cloned on a second plasmid (Paczkowski et al., 2017; Pearson, Pesci, & Iglewski, 1997). A set of HSLs differing in both carbon chain length and in functionality at the C-3 carbon were synthesized using a modification of a previously reported method (Chhabra et al., 2003) (see Appendix). Figure 1A shows reporter output following addition of 100 nM of the cognate autoinducer, 3OC_12_HSL, and four other HSLs (3OC_14_HSL, 3OC_10_HSL, 3OC_8_HSL, and 3OC_6_HSL). All five HSLs activated LasR, but to differing levels. 3OC_12_HSL, 3OC_14_HSL, and 3OC_10_HSL elicited maximal LasR activity, and 3OC_8_HSL and 3OC_6_HSL stimulated 7-fold and 18-fold less activity, respectively. We examined 8 other HSLs harboring different functionalities on the C-3 carbon in combination with the various tail lengths. Using dose-response analyses, we obtained EC_50_ values for the compounds (Table 1 and Table S1). 3OC_12_HSL was the most potent ligand, with an EC_50_ of approximately 2.8 nM. 3OC_14_HSL was half as potent with an EC_50_ of 5.6 nM. From there, the EC_50_ values followed the order 3OC_10_HSL < 3OC_8_HSL < 3OC_6_HSL. Table EV1 shows the remainder of the data and that the ketone versions of the molecules are the most potent for every chain length. For this reason, we used the five HSLs shown in Table 1 for much of the remainder of this work. We measured *in vivo* LasR activity in response to the test HSLs using an elastase assay (Gambello & Iglewski, 1991). Elastase is encoded by *lasB*, and as a reminder, we employed the *lasB* promoter in the *E. coli* reporter assay. For the elastase analyses, we used a Δ*lasI P. aeruginosa* strain that makes no endogenous 3OC_12_HSL. When supplied at 100 nM, all of the test compounds elicited some elastase activity, but 3OC_12_HSL stimulated the highest elastase production (Figure 1B). We do note that in *P. aeruginosa*, 3OC_14_HSL and 3OC_8_HSL stimulated lower activity than expected based on their EC_50_ values in *E. coli.* Indeed, as shown below, these two molecules had reduced activity in all assays in all *P. aeruginosa* strains used here. While we do not know the underlying molecular mechanism, we suspect that perhaps there is reduced permeability into *P. aeruginosa* and/or there is a factor in *P. aeruginosa* that does not exist in *E. coli* that binds and titrates out these two molecules. Nonetheless, our results indicate that, at least with respect to the HSLs we tested, *in vivo,* LasR is most active in response to its cognate autoinducer 3OC_12_HSL, but non-cognate HSLs can induce production of the quorum-sensing product elastase, and presumably other quorum-sensing regulated outputs.

**Figure 1.**
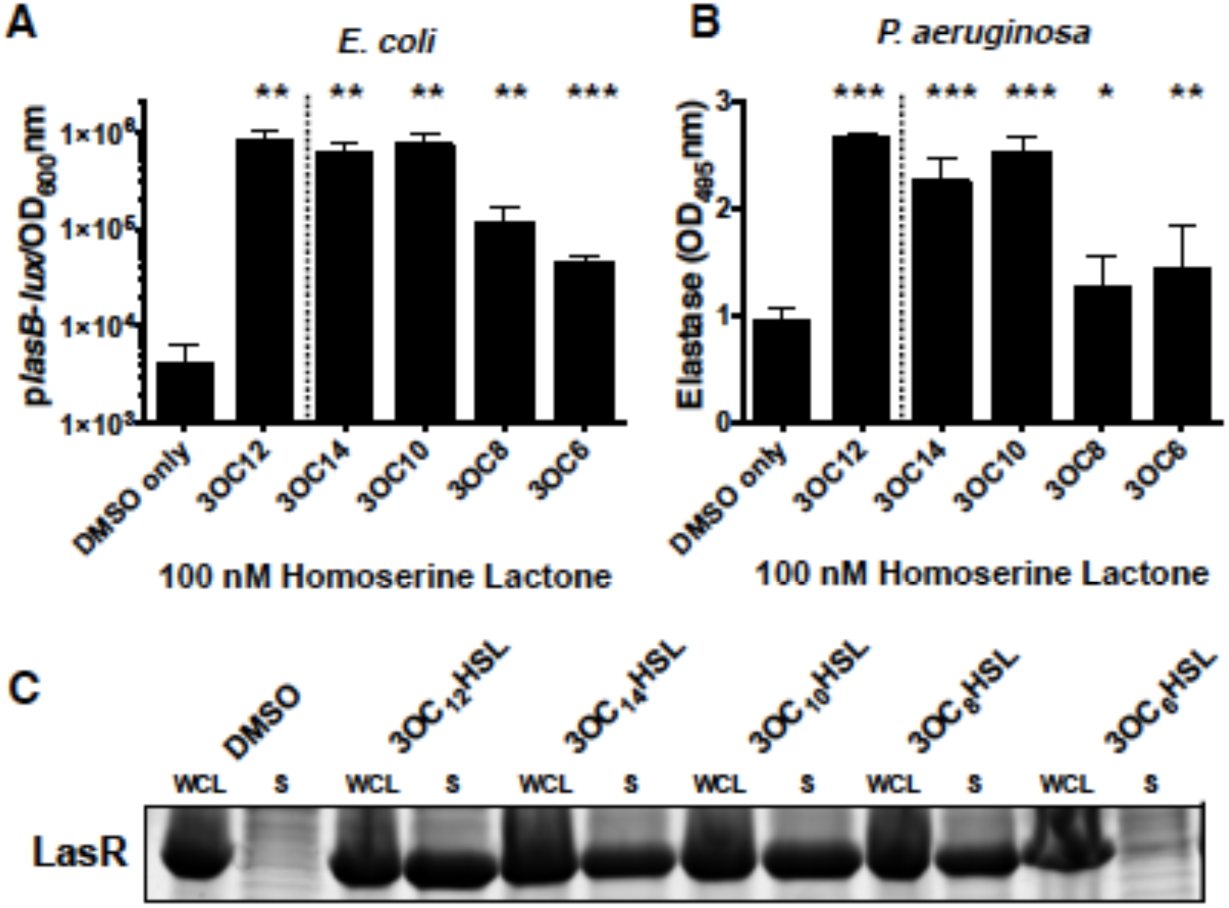
LasR is activated by multiple homoserine lactone autoinducers. A) LasR-dependent bioluminescence was measured in *E. coli*. Arabinose-inducible LasR was produced from one plasmid and the p*lasB-lux* reporter construct was carried on a second plasmid. 0.1% arabinose was used for LasR induction. B) Elastase activity was measured from Δ*lasI P. aeruginosa* using elastin-Congo red as the substrate. In A and B, 100 nM of the designated HSLs were tested. Two technical replicates were performed for each biological sample and 3 biological replicates were assessed. Error bars denote standard deviations of the mean. Paired 2-tailed t-tests were performed comparing each compound to the DMSO control. P-values: * <.05, ** <.01, *** <.0001. In panels A and B, we have separated the DMSO control and the results with the cognate autoinducer 3OC_12_HSL by the dotted vertical line. C) Comparison of LasR LBD protein levels in whole cell lysates (WCL) and in the soluble fractions (S) of *E. coli* cells that harbor the DNA encoding the LasR LBD on a plasmid. 1 mM IPTG was used for LasR LBD induction and either 1% DMSO or 10 μM of the indicated HSL was supplied. In all lanes, protein from .05 OD of cells was loaded. Results are representative of 3 trials.

**Table 1.** EC_50_ (nM) values for LasR and HSL compounds in the *plasB-lux* assay

**LasR Allele**
|  | WT | S129C | S129W | S129F | S129T | S129M | L130F |
| --- | --- | --- | --- | --- | --- | --- | --- |
| 3OC<sub>12</sub>HSL | 2.8 | 5.9 | 76.8 | 177 | 870 | 6570 | 2.0 |
| 3OC<sub>14</sub>HSL | 6.2 | 11.8 | 12.8 | 41.2 | 4070 | NR | 5.3 |
| 3OC<sub>10</sub>HSL | 8.0 | 31.5 | 2620 | 4980 | 4510 | NR | 5.0 |
| 3OC<sub>8</sub>HSL | 885 | 2860 | 2180 | NR | NR | NR | 80.7 |
| 3OC<sub>6</sub>HSL | 2370 | 4400 | NR | 55600 | 71700 | NR | 1390 |

NR denotes non-responsive

LasR and most other LuxR-type receptors fold around their cognate HSL ligands. Thus, they are only soluble, capable of dimerizing, and binding DNA when ligand is present (Bottomley, Muraglia, Bazzo, & Carfi, 2007; Zhu & Winans, 2001). The results in Figure 1A and 1B suggest that LasR can fold around HSLs in addition to 3OC_12_HSL. To verify this notion, we tested whether LasR could be solubilized by non-native HSLs. To do this, we grew *E. coli* producing the LasR LBD in the presence of the 13 HSL compounds (Figure 1C shows the five test compounds and Figure EV1A shows the eight other compounds in the collection). Consistent with previous results, in the absence of any ligand (DMSO control), the LasR LBD is present in the whole cell lysate, but not in the soluble fraction indicating that the protein is insoluble (O’Loughlin et al., 2013; Schuster, Urbanowski, & Greenberg, 2004). All five of the HSL test compounds except for 3OC_6_HSL solubilized the LasR LBD (Figure 1C). Nonetheless, we could only purify to homogeneity the LasR LBD bound to the ligands containing the longer acyl tails: 3OC_12_HSL, 3OC_14_HSL, and 3OC_10_HSL. Together, the results in Figure 1 suggest that although 3OC_8_HSL and 3OC_6_HSL can bind to and activate LasR, their interactions must be more transient than ligands with longer acyl tails. To garner evidence for this idea, we performed thermal shift analyses on LasR LBD-ligand complexes without and with the addition of either the same or a different HSL. The LasR LBD bound to 3OC_10_HSL, 3OC_12_HSL, and 3OC_14_HSL had melting temperatures of 42.3 °C, 49.1 °C, and 50.5 °C, respectively (Figure 2, black lines) showing that LasR stability increases with increasing ligand tail length. Notably, the LasR LBD is more stable when bound to the non-cognate ligand 3OC_14_HSL than when bound to the cognate ligand 3OC_12_HSL. The discrepancy between the enhanced stability of purified LasR LBD:3OC_14_HSL relative to 3OC_12_HSL in the thermal shift assay and the higher EC_50_ for 3OC_14_HSL compared to 3OC_12_HSL could result from increased hydrophobic interactions in the stably purified complex that do not drive activation and affinity of LasR for a particular ligand.

**Figure 2.**
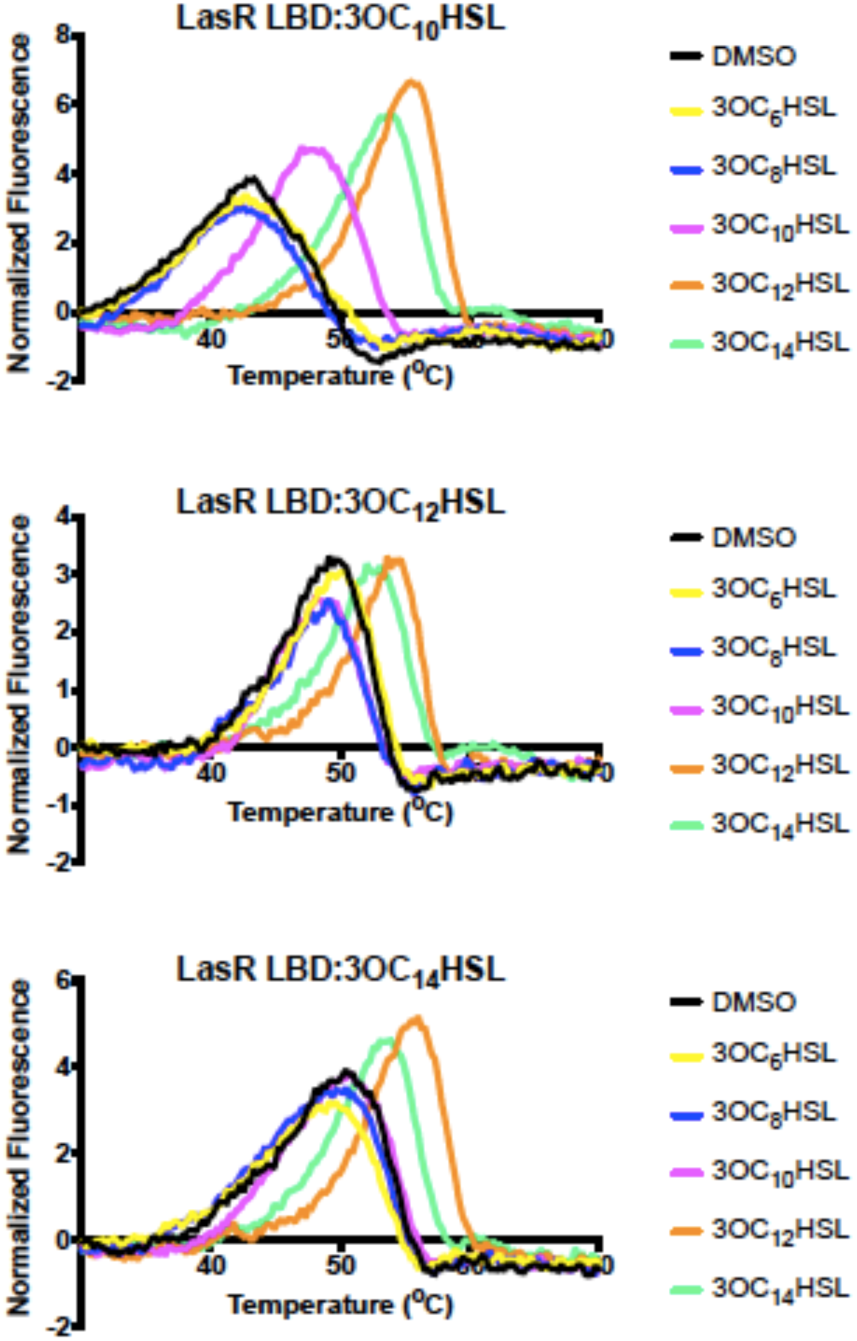
LasR is stable when bound to long chain HSLs. Thermal shift analyses of purified LasR LBD bound to 3OC_10_HSL (top), 3OC_12_HSL (middle), and 3OC_14_HSL (bottom) without (designated DMSO) and following supplementation with an additional 10 μM of the indicated HSLs. Each line represents the average of 3 replicates.

Exogenously supplied autoinducers can shift the melting temperature of an existing receptor-ligand complex if the added ligand has the ability to stabilize the unfolding protein as it releases the pre-bound ligand. Importantly, exogenously supplied HSLs can only stabilize the LasR LBD if they have affinities that are equal to or higher than the originally bound ligand (Paczkowski et al., 2017). In the case of the LasR LBD:3OC_10_HSL complex, when added exogenously, the ligands 3OC_10_HSL, 3OC_12_HSL, and 3OC_14_HSL stabilize the LasR LBD, increasing the melting temperature 3.4 °C, 6.4 °C, and 5.5 °C, respectively (Figure 2). By contrast, the LasR LBD:3OC_12_HSL and the LasR LBD:3OC_14_HSL complexes could only be further stabilized by exogenously supplied 3OC_12_HSL and 3OC_14_HSL, but not by 3OC_10_HSL, indicating that the association rate of 3OC_10_HSL for the LasR LBD is slower than that of 3OC_12_HSL and 3OC_14_HSL (Figure 2). While the short acyl chain HSLs were capable of activation of LasR with high micromolar EC_50_ values in the p*lasB-lux* reporter assay, 3OC_6_HSL and 3OC_8_HSL did not stabilize the LasR LBD protein sufficiently in *E. coli*, presumably due to their low affinities (Figure 1C and Table 1). Thus, they could not be tested in the traditional thermal shift assay. They also did not enhance the stabilization of the LasR LBD pre-bound with other ligands, analogous to the inability of 3OC_10_HSL to stabilize the LasR LBD:3OC_12_HSL complex as it melted (Figure 2). Together, our results indicate that in an environment containing a mixture of HSL autoinducers, LasR will preferentially detect long chain HSLs, with superior preference for its own autoinducer 3OC_12_HSL, followed closely by 3OC_14_HSL.

### LasR S129 drives ligand selectivity and sensitivity

To understand how LasR selects HSL ligands with which to interact, we performed site directed mutagenesis guided by the existing crystal structure of the LasR LBD bound to 3OC_12_HSL (Bottomley et al., 2007). We first focused on residue S129. Previous work has demonstrated that alteration of serine to alanine at this site enables some synthetic LasR activators to transform into inhibitors and vice-versa (Gerdt, McInnis, Schell, & Blackwell, 2015; Gerdt, McInnis, Schell, Rossi, & Blackwell, 2014). Moreover, the LasR LBD structure suggests that S129 is part of the network that interacts with the ligand acyl tail (Bottomley et al., 2007). For these reasons, we predicted that S129 could contribute to LasR selectivity. We constructed LasR S129C, S129W, S129F, S129T, and S129M and examined their activities in the *E. coli* p*lasB-lux* reporter system. Table 1 shows the EC_50_ values for all of the mutants and the five test compounds. The EC_50_ values follow the order: wildtype < LasR S129C < LasR S129W < LasR S129F < LasR S129T < LasR S129M. We note there are two exceptions to this trend among the low affinity interactions (Table 1). However, in those cases, the EC_50_ values are high and we suspect that the differences are not meaningful. The most severe mutation, LasR S129M responded exclusively to 3OC_12_HSL but it was the least sensitive of all the mutants to this ligand. Conversely, among the mutants, LasR S129C responded to the largest range of HSL varieties and it was the most sensitive. In Figure 3A, we provide the assay results for LasR S129F with the five test HSLs. We chose LasR S129F as the representative mutant for in-depth analysis because it had intermediate EC_50_ values among those obtained for this set of mutants. We assayed LasR S129F at 10 μM of each compound, because increased ligand concentration was required to activate LasR S129F compared to wildtype LasR (see Table 1 and Figure 1). At 10 μM ligand, wildtype LasR and LasR S129F maximally responded to 3OC_12_HSL and 3OC_14_HSL. However, compared to wildtype LasR, LasR S129F was 7-fold less responsive to 3OC_10_HSL, 13-fold less responsive to 3OC_6_HSL and showed almost no response to 3OC_8_HSL. In the Δ*lasI P. aeruginosa* strain (Figure 3B), LasR S129F displayed a similar pattern. Together, our data indicate that alteration of LasR S129 restricts the set of HSLs that can activate LasR but also increases the EC_50_ values of activating HSLs. We conclude that LasR S129 drives selectivity. While the LasR S129 variants improve selectivity, they diminish sensitivity. This finding suggests that a trade-off exists between selectivity and EC_50_.

**Figure 3.**
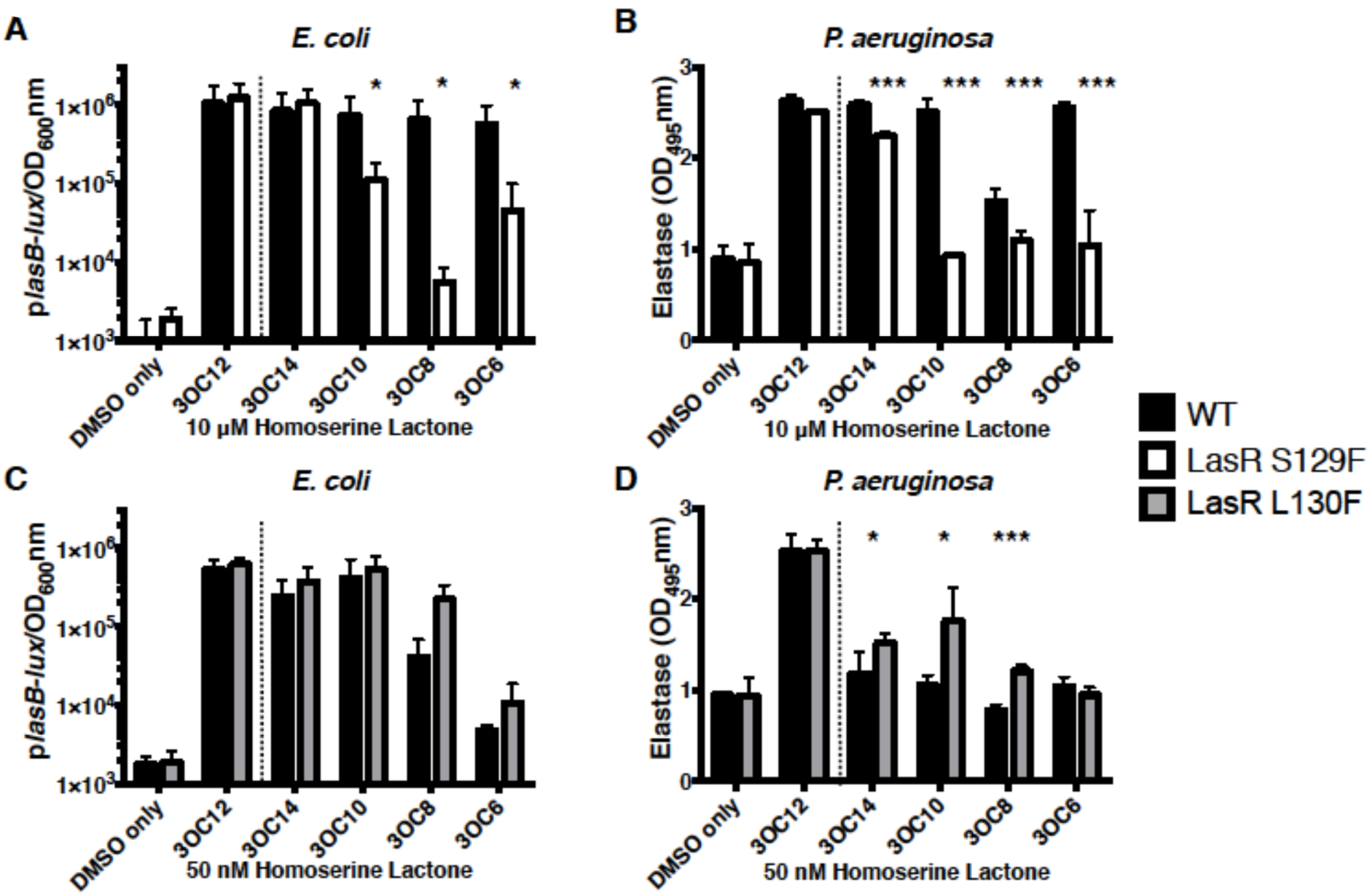
LasR S129F and LasR L130F alter LasR sensitivity to HSL autoinducers. A) Wildtype LasR and LasR S129F-driven bioluminescence from the *E. coli* p*lasB-lux* reporter (see Figure 1, panel A for detail). B) Wildtype LasR and LasR S129F-driven elastase activity in Δ*lasI P. aeruginosa* (see Figure 1, panel B for detail). In A and B, 10 μM of the indicated HSLs were provided. C and D) As in panels A and B, respectively, with wildtype LasR and LasR L130F and 50 nM of the indicated HSLs. Black bars, wildtype LasR; white bars, LasR S129F; gray bars, LasR L130F. Two technical replicates were performed for each biological sample and 3 biological replicates were assessed. Error bars denote standard deviations of the mean. Unpaired 2-tailed T-tests were performed comparing WT LasR to the mutant LasR for each compound. p-values: * < .05, *** <.001. In all panels, we have separated the DMSO control and the results with the cognate autoinducer 3OC_12_HSL by the dotted vertical line.

### LasR L130F drives ligand sensitivity

Because LasR S129F improves ligand selectivity and reduces sensitivity, we suspected that other residues in the vicinity could also affect ligand selectivity. For this reason, we exchanged L128 and L130 for phenylalanine. As above, we tested their responses to the set of representative HSLs using the *E. coli* p*lasB-lux* reporter assay. Table 1 shows that, surprisingly, LasR L130F exhibited a lower EC_50_ for every autoinducer tested. LasR L128F also conferred improved sensitivity to some short chain HSLs (Table S1), but its phenotype was less pronounced than that of LasR L130F, so we further analyzed the LasR L130F mutant here. Figure 3C shows the wildtype LasR and the LasR L130F responses to a low concentration (50 nM) of the test HSLs. At this concentration, LasR L130F was roughly equivalent to wildtype LasR with respect to the response to 3OC_12_HSL, 3OC_14_HSL, and 3OC_10_HSL. However, LasR L130F was approximately 5-fold more responsive to 3OC_8_HSL than wildtype LasR and it was modestly more responsive to 3OC_6_HSL. When we performed the *P. aeruginosa* elastase assay at 50 nM test compound (Figure 3D), wildtype LasR only responded to 3OC_12_HSL, whereas LasR L130F responded to 3OC_14_HSL, 3OC_10_HSL, and 3OC_8_HSL in addition to 3OC_12_HSL. Collectively, these results show that LasR L130F is more sensitive to HSLs than wildtype LasR but it is less selective.

We used thermal shift analyses to explore the mechanism underlying the increased sensitivity of the LasR L130F mutant for HSLs. We used 3OC_12_HSL and 3OC_14_HSL as our test ligands. Compared to the wildtype LasR LBD, the LasR LBD L130F is more stable when bound to each ligand (Figure 4A), suggesting that the L130F alteration increases the overall stability of the LasR protein. We exploited this feature to successfully purify the LasR LBD L130F bound to 3OC_8_HSL. As mentioned above, we could not purify the wildtype LasR LBD bound to 3OC_8_HSL. We performed thermal shift analyses on the LasR LBD L130F bound to 3OC_8_HSL, 3OC_10_HSL, 3OC_12_HSL, and 3OC_14_HSL to which we added different HSLs. Similar to the wildtype LasR LBD, exogenous addition of HSLs with long acyl chains further enhanced the stability of the LasR LBD L130F:HSL complexes compared to when HSLs with shorter acyl chains were added (Figure 2 and Figure 4B). For instance, the LasR LBD L130F:3OC_8_HSL stability was enhanced by the addition of 3OC_8_HSL, 3OC_10_HSL, 3OC_12_HSL, and 3OC_14_HSL but not 3OC_6_HSL (Figure 4B). Protein solubility analyses track with these findings; long acyl chain ligands solubilize the LasR LBD L130F protein whereas short chain ligands do not (Figure 4C, Figure S1B). Consistent with our EC_50_ values, chain length appears to be the most important factor driving protein solubility and stabilization for both the LasR LBD and the LasR LBD L130F (Figure S1B and Figure 4B, respectively).

**Figure 4.**
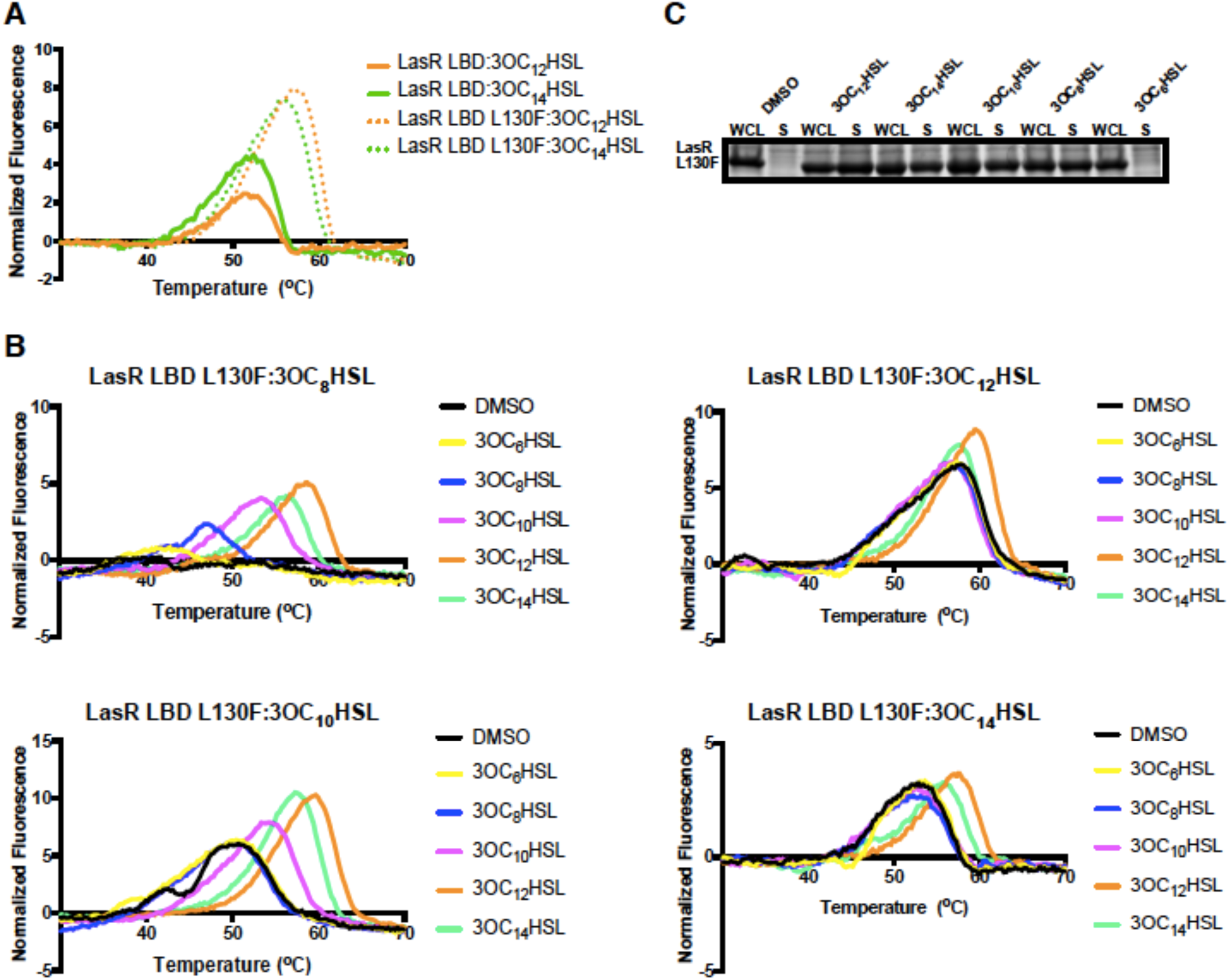
The LasR LBD L130F is more stable than the wildtype LasR LBD. A) Thermal shift analyses of purified LasR LBD (solid lines) and LasR LBD L130F (dotted lines) bound to 3OC_12_HSL (orange) or 3OC_14_HSL (green). Each line represents the average of 3 replicates. B) Thermal shift analyses of purified LasR LBD L130F bound to 3OC_8_HSL (top left), 3OC_10_HSL (bottom left), 3OC_12_HSL (top right), and 3OC_14_HSL (bottom right) without (designated DMSO) and following supplementation with an additional 10 μM of the indicated HSLs. C) Comparison of LasR L130F levels in the whole cell lysates (WCL) and the soluble fractions (S) of *E. coli* cells harboring the DNA encoding LasR LBD L130F on a plasmid (see Figure 1, panel C for details). Either 1% DMSO or 10 μM of the indicated HSL molecule was added.

### LasR ligand selectivity and sensitivity influence the timing of quorum-sensing-controlled traits

We have shown that LasR L130F detects and responds to HSLs at lower concentrations than does wildtype LasR, whereas LasR S129F requires higher concentrations. Thus, we predicted that introducing these alleles into *P. aeruginosa* should influence its quorum-sensing responses in opposing manners. To test this prediction, we assayed pyocyanin production over time as the quorum-sensing readout in response to 3OC_12_HSL, 3OC_14_HSL, and 3OC_8_HSL in a Δ*lasI P. aeruginosa* strain containing wildtype *lasR*, *lasR S129F*, or *lasR L130F*. Guided by the EC_50_ values for each HSL, we tested appropriate low and high concentrations of each HSL to set windows that would enable us to observe responses from the wildtype and each mutant. Consistent with their relative EC_50_ values, when a low concentration of 3OC_12_HSL (50 nM) was added, the strain with LasR L130F made more pyocyanin than the strain with wildtype LasR at every time point (Figure 5A). At this HSL concentration, the strain carrying LasR S129F never activated pyocyanin production, presumably because the concentration of 3OC_12_HSL was far below the EC_50_ for LasR S129F (Figure 5A). When the same assay was performed with 1μM 3OC_12_HSL, rapid and maximal pyocyanin output occurred for *P. aeruginosa* carrying both wildtype LasR and LasR L130F. Furthermore, this high ligand concentration reveals that LasR S129F can drive pyocyanin production in response to 3OC_12_HSL, albeit weakly and only after 5.5 hours (Figure 5B). A similar pattern was observed for the low concentration of 3OC_14_HSL. Specifically, LasR L130F activated pyocyanin production earlier than wildtype LasR, and LasR S129F failed to activate pyocyanin production (Figure 5C). At the high concentration of 3OC_14_HSL, LasR S129F did activate pyocyanin production, however later and less strongly than wildtype LasR and LasR L130F (Figure 5D). With respect to 3OC_8_HSL, at both low and high concentrations, all of the responses were considerably weaker than with the longer chain HSLs, nonetheless, in each case, LasR L130F more strongly activated pyocyanin production than wildtype LasR (Figure 5E, F). LasR S129F showed no response in the 3OC_8_HSL assays (Figure 5E, F).

**Figure 5.**
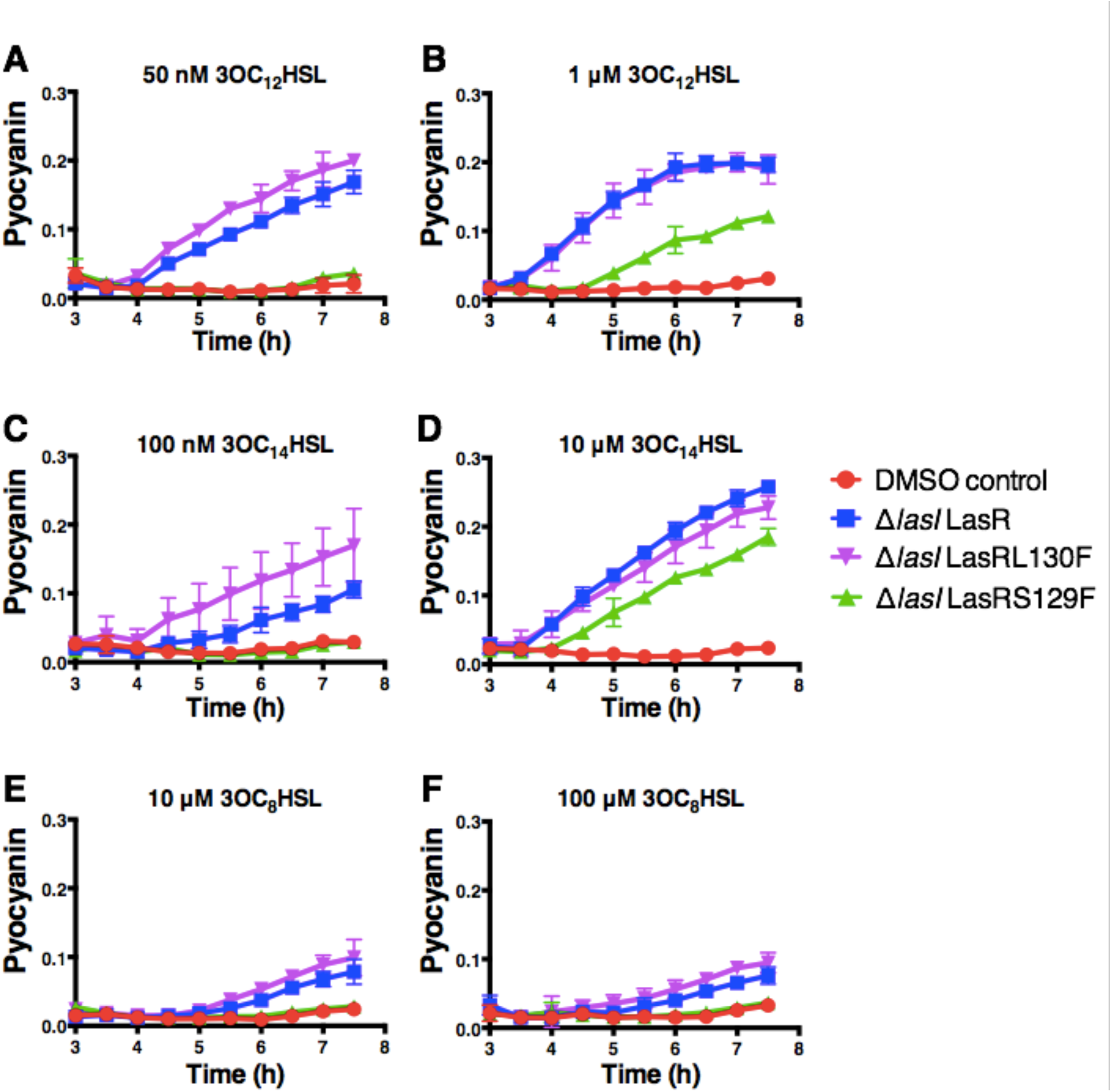
Wildtype LasR, LasR S129F, and LasR L130F display distinct pyocyanin production phenotypes in response to different HSL autoinducers. Pyocyanin production was measured spectophotometrically in Δ*lasI P. aeruginosa* over the growth curve. Y-axis “Pyocyanin” is the amount of pyocyanin pigment (OD_695_ nm) over cell density (OD_600_ nm). Designations are: red, DMSO control; blue, wildtype LasR; purple, LasR L130F; green, LasR S129F. Data show the mean of 3 biological replicates. Error bars denote standard deviations of the mean. Concentrations and ligands used are: A) 50 nM 3OC_12_HSL, B) 1 μM 3OC_12_HSL, C) 100 nM 3OC_14_HSL, D) 10 μM 3OC_14_HSL, E) 10 μM 3OC_8_HSL, and F) 100 μM 3OC_8_HSL

We note one curious finding with 10 μM 3OC_14_HSL in Figure 5D. At this concentration, wildtype LasR activated pyocyanin earlier and more strongly than LasR L130F, which does not track with the EC_50_ values and all of our above companion analyses. We propose that this phenotype is due to RsaL accumulation. RsaL is a quorum-sensing regulator whose expression is activated by LasR (Rampioni et al., 2007; Schuster & Greenberg, 2007). RsaL represses pyocyanin production genes (Cabeen, 2014; Schuster & Greenberg, 2007). We propose that because LasR L130F is more active than wildtype LasR, at high autoinducer concentration, LasR L130F possesses increased activity relative to wildtype LasR, and so LasR L130F stimulates higher RsaL production than does wildtype LasR. As a consequence, partial pyocyanin inhibition occurs. This logic suggests that, at high autoinducer concentration, wildtype LasR would activate sufficient RsaL production to cause pyocyanin inhibition. Indeed, addition of 100 μM 3OC_12_HSL to the Δ*lasI* strain carrying wildtype LasR induced less pyocyanin production than when 1 μM 3OC_12_HSL was added (Figure S2). We show that RsaL is responsible for this phenotype by deleting the *rsaL* gene. As expected, the Δ*lasI* Δ*rsaL* strain produced increased overall pyocyanin relative to the Δ*lasI* strain. Importantly, however, unlike the Δ*lasI* strain, the Δ*lasI* Δ*rsaL* double mutant does not produce more pyocyanin in response to 1 μM 3OC_12_HSL than in response to 100 μM 3OC_12_HSL (Figure S2). We never observed pyocyanin reductions in the strains carrying any of the LasR alleles when 3OC_8_HSL was added, even at 100 μM. We suspect this is because 3OC_8_HSL is such a poor agonist that there is not enough LasR activity at any concentration to stimulate high level RsaL accumulation.

### Structural basis underlying LasR ligand preferences

To determine the molecular basis enabling LasR to accommodate non-native autoinducers, we determined the structures of the LasR LBD L130F bound to 3OC_10_HSL and 3OC_14_HSL. As mentioned, the wildtype LasR LBD structure bound to the native autoinducer, 3OC_12_HSL already exists (Bottomley et al., 2007). Our rationale for using the LasR LBD L130F mutant for these studies was that, its inherently enhanced stability, as judged by the thermal shift data (Figure 4A), suggested that it would be ideal for crystallographic studies that are not possible with the wildtype LasR LBD. Indeed, we could determine the structures of LasR LBD L130F:3OC_10_HSL and LasR LBD L130F:3OC_14_HSL to 2.1 Å and 1.9 Å, respectively (Table A1). This resolution is sufficient to observe ligand occupancy and to compare with the LasR LBD:3OC_12_HSL structure (Figure 6A). The L130F residue is buried in a hydrophobic pocket distal to the ligand binding pocket (Figure S3A). Phenylalanine, rather than leucine at this position, provides increased hydrophobic interactions with amino acid residues L23, L30, F32, I35, L114, L118, L128, L151, and L154. We suggest that this arrangement increases the overall stability of the LasR protein, which in turn, allows it to accommodate an expanded set of HSLs compared to wildtype LasR.

**Figure 6.**
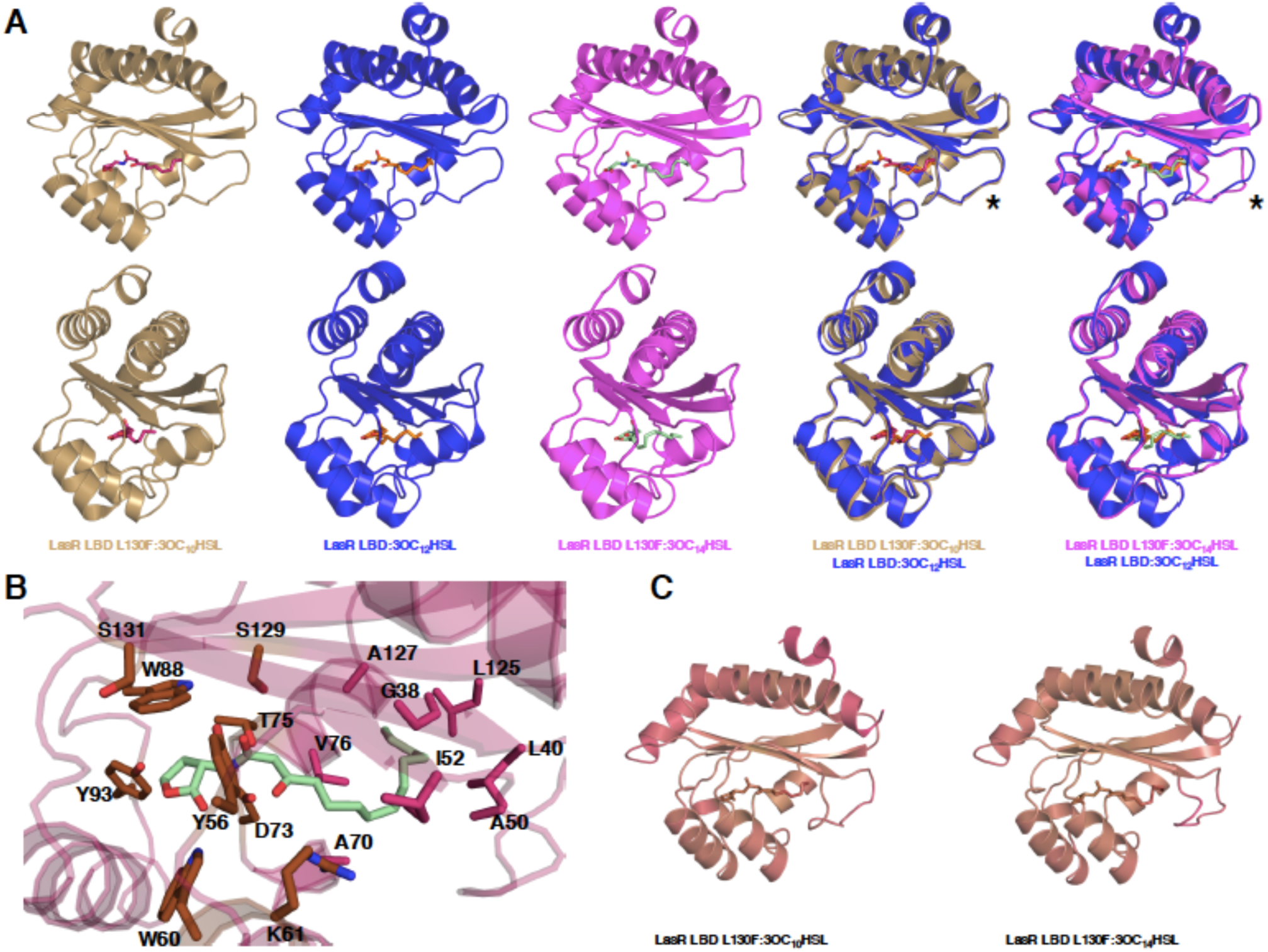
Crystal structures of LasR LBD L130F bound to 3OC_10_HSL and 3OC_14_HSL. A) Crystal structures of LasR LBD L130F:3OC_10_HSL (gold) and LasR LBD L130F:3OC_14_HSL (magenta) compared to the wildtype LasR LBD:3OC_12_HSL structure (blue, modified from Bottomley et al. 2007, PBD: 2UVO). The bottom images show 90-degree rotations of the crystal structures relative to the images above. In the top right-most 2 structures, the asterisks highlight the LasR loop region that includes residues 40-52 and that undergoes a conformational shift when 3OC_14_HSL is bound. B) LasR LBD L130F:3OC_14_HSL crystal structure (protein: magenta, ligand: green). Amino acids drawn in stick format show important residues for lactone head binding (Y56, W60, K61, D73, T75, W88, Y93, S129, and S131, all colored in brown) and acyl tail binding (G38, L40, A50, I52, A70, V76, L125, and A127, all colored in pink). C) Comparison of the average B-factors for LasR LBD L130F bound to 3OC_10_HSL and bound to 3OC_14_HSL. The structures are colored from gold to magenta with gold representing the lowest average B-factor and magenta representing the highest average B-factor.

To understand how LasR can accommodate multiple long chain HSL ligands in its binding site, we used the crystal structures to examine the key residues and regions of the binding pocket that interact with the different ligands. In all the structures, the lactone head groups have the exact same placement, likely due to strong hydrogen bonding between the lactone ring carbonyl moiety and residue W60 (Figure 6B, displayed in brown). The lactone head group and the ketone moiety on carbon 3 are further stabilized by an extensive hydrogen bonding network comprised of residues Y56, K61, D73, T75, W88, Y93, S129, and S131 (Figure 6B, displayed in brown). In the three structures, the C_10_, C_12_, and C_14_ tails take similar paths, until carbon 6, where C_10_ and C_14_ diverge from the path taken by the C_12_ tail in the original structure. Remarkably, this departure in tail path enables both the shorter and longer tails to occupy a similar hydrodynamic volume as the tail on 3OC_12_HSL (Figure S3B). The volume constraint is likely established by hydrophobic interactions with residues G38, L40, A50, I52, A70, V76, L125, and A127 (Figure 6B, displayed in pink). These same residues are also responsible for stabilizing 3OC_10_HSL in the ligand binding pocket. However, because the C_10_ tail is two carbons shorter than that of the native ligand, the terminal carbons in 3OC_10_HSL have higher intrinsic flexibility at the ligand-protein interface. The measured B-factors suggest increased flexibility stems from less stable hydrophobic interactions (Figure 6C).

We next investigated whether there were any structural rearrangements in the crystals that could account for the expanded HSL binding capabilities of LasR L130F compared to wildtype LasR. An ∼2 Å shift occurs in the loop corresponding to residues 40-52 (highlighted by asterisks in Figure 6A) in the LasR LBD L130F:3OC_14_HSL structure compared to the LasR LBD L130F protein complexed with 3OC_10_HSL and the wildtype LasR LBD complexed with 3OC_12_HSL (Figure 6A and 6C). This loop corresponds to a region with above average B-factor, as depicted by magenta coloring in Figure 6C. The alteration in the positioning of the loop indicates that it could be important for accommodating different HSLs. To test this possibility, we mutated LasR residue Y47 in this loop. We chose Y47 because it is the residue that shifts the most among the different structures. LasR Y47S and LasR Y47R displayed decreased sensitivity to both 3OC_12_HSL and 3OC_14_HSL in the *E*. *coli* reporter assay (Figure S4) suggesting that the loop provides interactions with the ligand tails that foster increased protein stability, allowing LasR to be activated.

To understand the structural basis underlying specificity and promiscuity in this family of proteins, we compared the structures of SdiA LBD:3OC_8_HSL, CviR LBD:C_6_HSL, and TraR LBD:3OC_8_HSL to LasR LBD L130F:3OC_14_HSL (Figure 7A). We chose these particular structures because the receptors display a range of ligand selection preferences – from strict to promiscuous. Similar to what we show here for LasR, SdiA is promiscuous with respect to ligand selectivity (Michael et al., 2001; Nguyen et al., 2015; Sitnikov et al., 1996). Conversely, CviR and TraR display strict specificity for their native autoinducers (Chen et al., 2011; Vannini et al., 2002; Zhang et al., 2002; Zhu & Winans, 2001). In terms of overall tertiary structure, a loop similar to the one we pinpoint in Figure 6A as critical for LasR to accommodate different ligands, exists in the SdiA LBD (Figure 7A). No such loop exists in the CviR and TraR LBD structures (Figure 7A). This result is consistent with the idea that this flexible loop confers ligand promiscuity.

**Figure 7.**
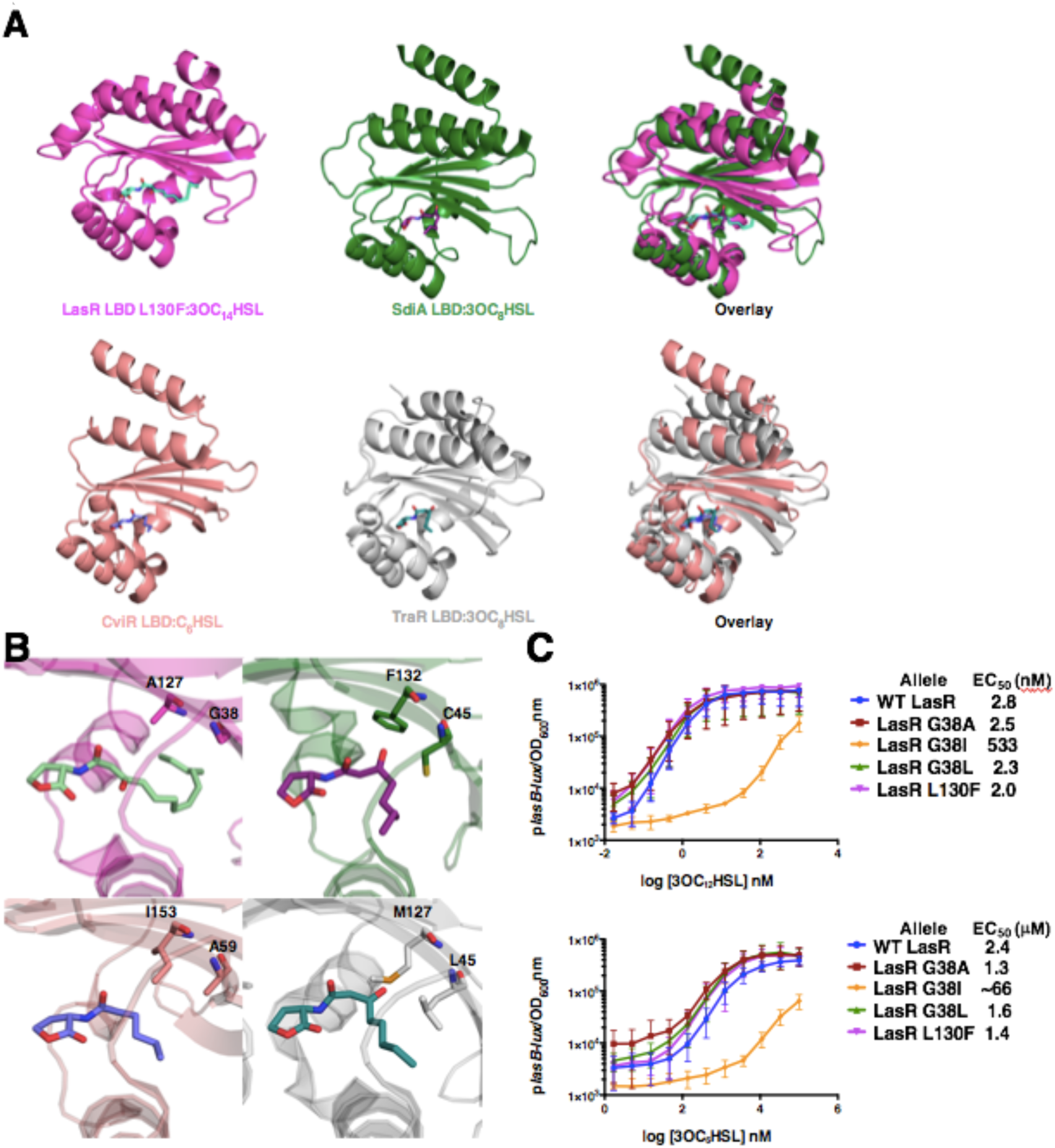
A conserved flexible loop region confers promiscuity to LuxR-type receptors. A) Top images: structural comparison of the LuxR-type receptors LasR LBD L130F:3OC_14_HSL (magenta) and SdiA LBD:3OC_8_HSL (green, modified from Nguyen et al., 2015, PDB: AY17) that exhibit promiscuity with respect to ligand binding. Bottom images: structural comparison of the LuxR-type receptors CviR LBD:C_6_HSL (pink, modified from Chen et al., 2011, PDB: 3QP1) and TraR LBD:3OC_8_HSL (silver, modified from Zhang et al., 2002, PDB: 1L3L) that display strict ligand specificity. B) Structural comparison of the protein:ligand interfaces for LasR LBD L130F:3OC_14_HSL (top left, magenta), SdiA LBD:3OC_8_HSL (top right, green), CviR LBD:C_6_HSL (bottom left, pink), and TraR LBD:3OC_8_HSL (bottom right, silver). Residues that make important hydrophobic sidechain interactions in each protein:ligand complex are shown in stick format and named. C) Dose response curves from the *E. coli* p*lasB-lux* reporter assay (see Figure 1, panel A) were used to determine EC_50_ values for 3OC_12_HSL (top panel) and 3OC_6_HSL (bottom panel) for wildtype LasR (blue), LasR G38A (dark red), LasR G38I (orange), LasR G38L (dark green), and LasR L130F (magenta). Two technical replicates were performed for each biological sample and 3 biological replicates were assessed. Error bars denote standard deviations of the mean.

In addition to the differences in the overall structures of the receptors, we noted different conformations for the acyl chains in the LasR LBD structures compared to those in the other LuxR-type receptors. The autoinducer acyl chains in the CviR, TraR, and SdiA LBD structures have similar conformations within the ligand binding pockets terminating between residues Y88 and M89 for CviR, Y61 and F62 for TraR, and Y71 and Q72 for SdiA. By contrast, the acyl chains of the different ligands in the LasR LBD L130F structures orient their terminal carbons toward the opposing face of the ligand binding pocket (Figure 7B). These different paths appear to be driven by the hydrophobic interactions we described above for the different LasR ligands (Figure 6B). We identified eight hydrophobic residues (G38, L40, A50, I52, A70, V76, L126, and A127) that dictate the shape of the ligand binding pocket in LasR. Indeed, these residues have generally hydrophobic characteristics in all LuxR-type proteins, but the size of each sidechain varies (Figure 7B). For example, A127 in LasR corresponds to F132 in SdiA. The smaller residue in LasR accommodates the altered path taken by its autoinducer, allowing the terminal carbon of the acyl chain to bind proximal to residue A127. A bulkier residue at this position, as in SdiA, sterically hinders this path for the ligand, forcing the acyl chain to bend in the opposite direction. Thus, in SdiA, F132 forces the terminal carbon of 3OC_8_HSL to bind distally. Indeed, the I153 residue in CviR and the M127 residue in TraR appear to perform roles analogous to F132 in SdiA in dictating the distal orientations of their ligands.

Our crystal structures suggest that LasR G38 could also be important for allowing HSL tails to adopt a conformation different from those in SdiA, TraR, and CviR and, in so doing, possibly affect LasR ligand selection (Figure 7B). SdiA, TraR, and CviR all contain larger amino acid sidechains (C45, L45, and A59, respectively) at this position (Figure 7B). LasR G38 lies proximal to carbons 9 and 10 in the acyl chains of all the ligands in all the LasR LBD structures. Presumably, a bulkier side chain would sterically hinder the preferred tail paths for 3OC_12_HSL or 3OC_14_HSL, but might enhance the binding of short chain HSLs, such as 3OC_6_HSL, due to increased hydrophobic interactions. To test this prediction, we mutated G38 in LasR to bulkier amino acids: leucine, isoleucine, and alanine. We tested these alleles in our *E. coli* p*lasB-lux* reporter in response to both a short (3OC_6_HSL) and a long (3OC_12_HSL) chain HSL (Figure 7C). LasR G38I produced low p*lasB-lux* activity with both molecules suggesting that isoleucine is too bulky to accommodate any HSL well. Consistent with this interpretation, the EC_50_ values for LasR G38I were 533 nM for 3OC_12_HSL and ∼66 μM for 3OC_6_HSL. By contrast, LasR G38A and LasR G38L fully activated the reporter in response to both test ligands (Figure 7C). The EC_50_ values for wildtype LasR, LasR G38A, LasR G38L, and LasR L130F were equivalent for 3OC_12_HSL (2.8 nM, 2.5 nM, 2.3 nM, and 2.0 nM, respectively, Figure 7C, top panel). However, the EC_50_ values for LasR G38A and LasR G38L were lower (1.3 μM and 1.6 μM, respectively) than that of wildtype LasR (2.4 μM) for 3OC_6_HSL (Figure 7C, bottom panel). These values are comparable to that of LasR L130F for 3OC_6_HSL (1.4 μM, Figure 7C, bottom panel). Given that we have established that LasR L130F is more stable than wildtype LasR, irrespective of ligand chain length, we suggest that the findings in Figure 7 support the hypothesis that larger amino acid residues at G38 improve the LasR affinity for shorter chain HSLs by forming more stable protein-ligand complexes.

## Discussion

*P. aeruginosa* can live in environments in which it encounters other bacterial species that produce HSL autoinducers (Tashiro, Yawata, Toyofuku, Uchiyama, & Nomura, 2013). Our results show that LasR can, with reduced affinity, detect non-native HSLs and in response, activate transcription of quorum-sensing target genes. We suggest that determining the mechanisms that promote or restrict ligand access to LasR is important for understanding how the *P. aeruginosa* quorum-sensing response could be naturally or synthetically manipulated. Here, we identified mutations that alter the selectivity and sensitivity of LasR. We found that LasR L130F improves LasR sensitivity, but at the cost of decreased selectivity. In contrast, we find that alterations at LasR S129 can increase selectivity for autoinducer recognition but, that feature is offset by a reduction in sensitivity. Our findings suggest that LasR balances ligand sensitivity with selectivity. The LasR S129 mutants show that decreasing the LasR sensitivity to ligands dampens and delays the quorum-sensing response (Figure 3B and Figure 5). We presume that inadequate activation decreases the potency of *P. aeruginosa* virulence. By contrast, the more sensitive LasR L130F allele causes higher and premature activation of quorum sensing (Figure 3D and Figure 5). In this case, the release of expensive public goods before the population is at a sufficient cell density to optimally use those goods could promote cheating. LasR L130F may also negatively influence virulence at high autoinducer concentration by stimulating overproduction of RsaL (Figure 5D).

We investigated the published sequences of hundreds of clinical isolates of *P. aeruginosa* and we did not find any strain that possessed a mutation at either S129 or L130 (Feltner et al., 2016). We conclude that it is detrimental for LasR to become either more sensitive to or less selective for ligands. Perhaps, striking the ideal balance between selectivity and sensitivity in LasR means that detection of some non-native HSLs must be tolerated, although, importantly, only at higher concentrations relative to the native autoinducer.

We propose that LasR promiscuity could serve an important function in the environment (i.e., soil) and in eukaryotic hosts where *P. aeruginosa* encounters other species of bacteria. We suggest that LasR detection of non-native HSLs enables it to “eavesdrop” on its competitors. However, our finding that 3OC_12_HSL is the most potent agonist indicates that *P. aeruginosa* prioritizes detecting its own autoinducer over those of other species. Amongst our set of test compounds, 3OC_14_HSL is the second best LasR agonist and it is also highly potent, indicating that the LasR receptor has not optimized against detection of longer chain HSLs. Possibly, *P. aeruginosa* readily interacts with 3OC_14_HSL-producing bacteria. Alternatively, *P. aeruginosa* might not encounter 3OC_14_HSL in its natural surroundings as, presently, scant evidence exists for natural production of 3OC_14_HSL by bacteria. There are two preliminary reports of soil-dwelling nitrogen-fixing bacteria that are capable of making 3OC_14_HSL, 3OHC_14_HSL and/or C_14_HSL (Gao, Ma, Zhuang, & Zhuang, 2014; Mellbye, Spieck, Bottomley, & Sayavedra-Soto, 2017), but if, when, and how much they do remains mysterious. Specifically, mass spectrometry and bioassay show that *Nitrosospira briensis* produces 3OHC_14_HSL, but only at concentrations of 1 nM (Mellbye et al., 2017), too low for detection by wildtype LasR. Second, *Nitrospira multiformis* has a LuxI-type synthase (NmuI) that produces 3OC_14_HSL and C_14_HSL in recombinant *E. coli*. However, neither of these molecules solubilized the putative receptor, NmuR, and no conditions were identified under which *Nitrospira multiformis* produced the molecules (Gao et al., 2014). Future study will be necessary to determine the prevalence of HSLs longer than C_12_ in nature and whether *P. aeruginosa* encounters such molecules.

With the one exception of 3OC_14_HSL, wildtype LasR detects 3OC_12_HSL far more efficiently than other HSLs. Therefore, it is likely that, in mixed-species consortia, 3OC_12_HSL out-competes all other autoinducers. This finding suggests that while LasR is capable of detecting non-cognate autoinducers, it does not do so when *P. aerugino*sa is the majority species. However, we propose that there could exist conditions under which low-affinity “eavesdropping” would benefit *P. aeruginosa.* First, when *P. aeruginosa* is at low cell density, provision of non-cognate ligands made by other bacterial species that are at higher cell density could induce *P. aeruginosa* to activate its quorum-sensing behaviors prematurely relative to when it is at low cell density in mono-species culture. If so, *P. aeruginosa* could synthesize toxic defensive products such as pyocyanin and rhamnolipids that endow it with an advantage over competing species (Baron & Rowe, 1981; Smalley, An, Parsek, Chandler, & Dandekar, 2015). Beyond defensive products, perhaps some of the quorum-sensing-controlled products produced under such conditions can be used exclusively by *P. aeruginosa* and not by competing species. If so, LasR promiscuity could grant *P. aeruginosa* a “last ditch” opportunity to survive in environments in which it is vastly outnumbered by competing species.

Our combined genetic, biochemical, and structural work revealed the molecular basis for non-native autoinducer recognition by LasR. There exists a key flexible loop that, if present (LasR, SdiA), endows particular LuxR-type receptors with the ability to bind to multiple HSL ligands, but if absent (TraR, CviR), the LuxR-type receptor is highly specific for a particular HSL ligand. Indeed, this structure-function analysis could explain why a competitive inhibitor of CviR, chlorolactone (CL), behaves as an agonist in *P. aeruginosa* (O’Loughlin et al., 2013). These findings are particularly enlightening when considering attempts to design inhibitors that specifically target different LuxR-type receptors. The flexible loop and hydrophobic residues in LasR near the acyl chain binding site that we pinpoint here will need to be taken into account when developing small molecule inhibitors that target LasR or other LuxR-type proteins that possess this feature. Designing molecules that target the flexibility of the loop region and/or that destabilize the protein could be explored for promiscuous receptors, such as LasR. By contrast, targeted screening around molecules that resemble CL could yield inhibitors for LuxR-type proteins that display strict specificity for their cognate autoinducers, such as CviR.

## Materials and Methods

### Site directed mutagenesis

Mutations in *lasR* were constructed on the pBAD-A-*lasR* plasmid (Paczkowski et al., 2017). Primers were designed using the Agilent Quikchange primer design tool and PCR with pFUltra polymerase (Agilent). PCR reactions were treated with DpnI to eliminate parental plasmid DNA and the plasmids with the mutant *lasR* genes were transformed into One Shot TOP10 chemically competent *E. coli* cells (Invitrogen). Reactions were plated on LB agar plates containing ampicillin (50 μg/mL) and individual mutants were verified via sequencing with primers for the *lasR* gene (ARM203 and ARM204). Primers and strains used in this work are listed in Table A2 and Table A3, respectively.

### *P. aeruginosa* strain construction

In-frame, marker-less *lasR* mutations were engineered onto the chromosome of *P. aeruginosa* PA14 using pEXG2-suicide constructs with gentamicin selection and *sacB* counter selection (Borlee, Geske, Blackwell, & Handelsman, 2010; Kukavica-Ibrulj et al., 2008). The *lasR* gene and 500 bp of flanking regions were cloned into pUCP18 (Schweizer, 1991). Site directed mutagenesis was performed as described above to construct point mutations in plasmid-borne *lasR*. The DNA carrying the mutant *lasR* genes was obtained from pUCP18 by restriction enzyme digestion with BamHI and EcoRI (NEB), and subsequently, the fragments were ligated into pEXG2. The recombinant pEXG2 plasmids were transformed into *E. coli* SM10λ*pir* and, from there, the plasmids were mobilized into *P. aeruginosa* PA14 via biparental mating (Mukherjee, Moustafa, Smith, Goldberg, & Bassler, 2017; Simon, Priefer, & Puhler, 1983). Exconjugants were selected on LB plates containing 30 μg/mL gentamicin and 100 μg/mL irgasan after overnight growth at 37° C. After recovery, 5% sucrose was used to select for loss of the plasmid. Candidate mutants were patched onto LB plates and LB plates containing 30 μg/mL gentamicin to select against the resistance marker. Colony PCR was performed on gentamicin sensitive patches with primers that annealed 500 bp [base pairs] upstream and downstream of *lasR* (ARM455 and ARM456). These PCR products were sequenced with *lasR* forward and reverse primers (ARM203 and ARM204).

### *E. coli* p*lasB-lux* reporter assay for LasR activity

The development of an assay that reports on LasR activity in response to exogenous ligands using luciferase as the readout has been described previously (Paczkowski et al., 2017). In brief, 2 μL of overnight cultures containing p*lasB-luxCDABE* and pBAD-A with either wildtype *lasR* or mutant *lasR* alleles were back diluted into 200 μL LB medium and placed into clear-bottom 96-well plates (Corning). The plates were shaken at 30° C for 4 h and 0.1% arabinose was added to each well along with a test HSL at the concentrations designated in the text and figures. To perform dose response analyses, 1 mM of each HSL was serially diluted 3-fold 10 times, and 2 μL of each dilution was added to the wells. Higher or lower HSL concentrations were assayed when EC_50_ values did not fall into this range. Plates were shaken at 30°C for 4 h. Bioluminescence and OD_600_ were measured using a Perkin Elmer Envision Multimode plate reader. Relative light units were calculated by dividing the bioluminescence measurement by the OD_600_ nm measurement. Non-linear regression was performed in Graphpad Prism6 to obtain EC_50_ values.

### Elastase assay

The *P. aeruginosa* PA14 Δ*lasI* strains carrying either wildtype or mutant *lasR* genes were grown overnight with shaking at 37° C in LB medium. Cultures were back diluted 1:50 in 3 mL of LB and grown for an additional 8 h with shaking at 37° C. Strains were back diluted 1:1000 into 3 mL of LB medium and test HSLs or an equivalent volume of DMSO were added to each culture. These cultures were grown overnight at 37° C with shaking. 1 mL of each culture was removed and the cells were pelleted by centrifugation at 16,100 x g. The supernatant was removed and filtered through a .22 μm filter (Millipore) and 100 μL of supernatant was added to 900 μL of 10 mM Na_2_HPO_4_ containing 10 mg of elastin-Congo red substrate (Sigma-Aldrich). These preparations were incubated at 37° C for 2 h. The mixtures were subjected to centrifugation at 16,100 x g for 10 min. The resulting supernatants were removed and OD_495_ nm measured with a Beckman Coulter DU730 spectrophotometer against a blank of H_2_O.

### Thermal shift assay

Thermal shift analyses of 6xHis-LasR LBD and 6xHisLasR L130F LBD bound to HSLs were performed as previously described (Paczkowski et al., 2017). In short, ligand-bound protein was diluted to 5 μM in reaction buffer (20 mM Tris-HCL pH 8, 200 mM NaCl, and 1 mM DTT [dithiothreitol]) containing DMSO or 10 μM HSL test compound in 18 μL total volume. The mixtures were incubated at room temperature for 15 min. 5000x SYPRO Orange (Thermo-Fisher) in DMSO was diluted to 200x in reaction buffer and used at 20x final concentration. 2 μL of 200x SYPRO Orange was added to the 18 μL sample immediately prior to analysis. Samples were subjected to a linear heat gradient of 0.05 °C/s, from 25 °C to 99 °C in a Quant Studio 6 Flex System (Applied Biosystems) using the melting curve setting. Fluorescence was measured using the ROX reporter setting.

### Pyocyanin time course

Overnight cultures of the *P. aeruginosa* Δ*lasI*, Δ*lasI lasR S129F*, Δ*lasI lasR L130F*, and Δ*lasI* Δ*rsaL* strains were grown in LB medium with shaking at 37° C. 1.5 mL of each culture was diluted into 50 mL of fresh LB medium. HSLs were added at the concentrations described in the text and figures and the cultures were shaken at 37° C for an additional 3 h. From there forward, 1 mL aliquots were removed every 30 min for 300 min and cell density (OD_600_ nm) was measured immediately using a Beckman Coulter DU730 Spectrophotometer. The aliquots were subjected to centrifugation at 16,100 x g for 2 min and the clarified supernatants were removed. The OD_695_ nm of the supernatants were measured. Pyocyanin activity was determined by plotting the OD_695_ nm/OD_600_ nm over time for each strain.

### Protein production, purification, and crystallography

Recombinant 6xHis-LasR LBD and 6xHis-LasR LBD L130F proteins bound to 3OC_8_HSL, 3OC_10_HSL, 3OC_12_HSL, or 3OC_14_HSL were purified as previously described for LasR LBD:3OC_12_HSL using Ni-NTA affinity columns followed by size exclusion chromatography (Paczkowski et al., 2017). 6xHis-LasR LBD bound to 3OC_10_HSL and 6xHis-LasR LBD bound to 3OC_14_HSL were crystallized by the hanging drop diffusion method. Diffraction data were processed using the HKL-3000 software package (Minor, Cymborowski, Otwinowski, & Chruszcz, 2006). The structures were solved using Phaser in Phenix (Adams et al., 2011; Afonine et al., 2012) by molecular replacement, with the structure of LasR LBD:3OC_12_HSL used as the search model (Bottomley et al., 2007). Model building was performed using Coot (Emsley & Cowtan, 2004; Emsley, Lohkamp, Scott, & Cowtan, 2010) and further refinement was accomplished using Phenix (Adams et al., 2011). Images of the structures were generated with PyMOL (DeLano, 2009). When we describe specific amino acid or amino acid-ligand interactions, we provide the image for the best resolved example in the asymmetric unit.

### Protein solubility assay

*E. coli* BL21 DE3 (Invitrogen) containing plasmid-borne 6xHis-LasR LBD or 6xHis-LasR LBD L130F were grown overnight and back diluted 1:500 in 20 mL of LB medium containing ampicillin (100 μg/mL). Cultures were grown to OD_600_ of 0.5 and protein production was induced with 1 mM IPTG [Isopropyl β-D-1-thiogalactopyranoside]. Upon addition of IPTG, the desired test HSL was also added at a final concentration of 10 μM, and the cultures were incubated at 25 °C with shaking for 4 h. Cells were harvested at 3,000 x g and resuspended in lysis buffer (500 mM NaCl, 20 mM Tris-HCL pH 8, 20 mM imidazole, 5% glycerol, 1 mM EDTA, and 1 mM DTT). The cells were lysed using sonication (1 s pulses for 15 s with a 50% duty cycle). The fraction we call the whole cell lysate was harvested after sonication. The soluble fraction was isolated by centrifugation at 32,000 x g. Samples were subjected to electrophoresis on SDS-PAGE gels (Biorad) and imaged with an Image Quant LAS4000 gel dock using the trans-illumination setting (GE Healthcare).

### Homoserine lactone synthesis

Unless otherwise indicated, all temperatures are expressed in °C (degrees Centigrade). All reactions were conducted at room temperature unless otherwise noted. ^1^H-NMR spectra were recorded on a Varian VXR-400, or a Varian Unity-400 at 400MHz [megahertz] field strength. Chemical shifts are expressed in parts per million (ppm, δ units). Coupling constants (*J*) are in units of hertz (Hz). Splitting patterns describe apparent multiplicities and are designated as s (singlet), d (doublet), t (triplet), q (quartet), m (multiplet), quin (quintet) or br (broad). Mass spectrometry analyses were performed on a Sciex API 100 using electrospray ionization (ESI). LCMS was carried out using a C-18 reverse phase column (2.1 ID, 3.5 micron, 50 mm). The column conditions were 98% water with 0.05%TFA and 2% MeOH [methanol] to 100% MeOH over 5.5 min. Analytical thin layer chromatography was used to verify the purity as well as to follow the progress of reaction(s). Unless otherwise indicated, all final products were at least 95% pure as judged by HPLC / MS. Synthesis is diagrammed in Appendix Figure 1.

#### General procedure for the synthesis of homoserine lactones

To a solution of (3*S*)-3-aminotetrahydrofuran-2-one (1.00 eq, HBr salt) and Et_3_N (3.00 eq) in DCM was added a solution of the acid chloride (1.00 eq) in DCM. The resulting reaction mixture was stirred at room temperature for 3 h. The reaction mixture was then diluted with H_2_O (5 mL), and extracted with DCM (3 x 5 mL). The organic layers were combined, washed with brine (10 mL), dried over Na_2_SO_4_, filtered, and concentrated under reduced pressure to give a residue. The residue was purified by silica gel column chromatography to give the desired homoserine lactone as a white solid.

#### (*S*)-*N*-(2-oxotetrahydrofuran-3-yl)hexanamide (BB0189)

Gradient elution with Petroleum ether/EtOAc = 3/1 to 1/1 afforded the title compound (425 mg, 96% yield, 98% purity by LC/MS) as a white solid. ^1^H NMR (CDCl_3_) δ = 6.26 (s, 1H), 4.63 (m, 1H), 4.50 (td, *J* = 1.0, 5.6, 1H), 4.33 (m, 1H), 2.87 (m, 1H), 2.27 (t, *J* = 7.4, 2H), 2.20 (m, 1H), 1.71 (t, *J* = 7.4, 2H), 1.34 (m, 4H), 0.90 (t, *J* = 7.1, 3H); MS (ESI) calculated for C_10_H_17_NO_3_: m/z = 199; found: m/z = 200 (M+H).

#### (S)-*N*-(2-oxotetrahydrofuran-3-yl)octanamide (BB0192)

Gradient elution with Petroleum ether/EtOAc = 3/1 to 1/1 afforded the title compound (270 mg, 96% yield, 99% purity by LC/MS) as a white solid. ^1^H NMR (CDCl_3_) δ = 5.94 (s, br, 1H), 4.54 (m, 2H), 4.29 (ddd, *J* = 5.9, 9.5, 11.3, 1H), 2.88 (m, 1H), 2.26 (t, *J* = 7.7, 2H), 2.13 (m, 1H), 1.63 (m, 2H), 1.30 (m, 8H), 0.89 (t, *J* = 6.6, 3H); MS (ESI) calculated for C_12_H_21_NO_3_: m/z = 227; found: m/z = 228 (M+H).

#### (*S*)-*N*-(2-oxotetrahydrofuran-3-yl)decanamide (BB0195)

Gradient elution with Petroleum ether/EtOAc = 10/1 to 1/1 afforded the title compound (632 mg, 94% yield, 99% purity by LC/MS) as a white solid. ^1^H NMR (CDCl_3_) δ = 6.01 (s, br, 1H), 4.55 (ddd, *J* =5.7, 8.6, 11.6, 1H), 4.47 (t, *J* = 9.2, 1H), 4.29 (ddd, *J* = 6.1, 9.4, 11.2, 1H), 2.87 (m, 1H), 2.25 (t, *J* = 7.7, 2H), 2.12 (m, 1H), 1.66 (m, 2H), 1.30 (m, 12H), 0.88 (t, *J* = 6.8, 3H); MS (ESI) calculated for C_14_H_25_NO_3_: m/z = 255; found: m/z = 256 (M+H).

#### (*S*)-*N*-(2-oxotetrahydrofuran-3-yl)dodecanamide (BB0198)

Gradient elution with Petroleum ether/EtOAc = 10/1 to 1/1 afforded the title compound (632 mg, 94% yield, 99% purity by LC/MS) as a white solid. ^1^H NMR (CDCl_3_) δ = 5.94 (s, br, 1H), 4.60-4.45 (m, 2H), 4.29 (m, 1H), 2.90 (m, 1H), 2.25 (t, *J* = 7.2, 2H), 2.14 (m, 1H), 1.64 (m, 2H), 1.35-1.22 (m, 16H), 0.88 (t, *J* = 6.7, 3H); MS (ESI) calculated for C_16_H_29_NO_3_: m/z = 283; found: m/z = 284 (M+H).

#### General Procedures for the synthesis of 3-oxo homoserine lactones

##### Procedure A

To a stirring solution of 2,2-dimethyl-1,3-dioxane-4,6-dione (Meldrum’s acid) (1.00 eq) and DMAP (2.00 eq) in DCM at 0°C was added a solution of the acid chloride (1.00 eq) in DCM. The resulting reaction mixture was allowed to warm to room temperature and stirred for 12 h. The reaction mixture was diluted with DCM (30 mL) and washed with cold 2N HCl (3 x 30 mL). The organic layer was separated, dried over Na_2_SO_4_, filtered and concentrated under reduced pressure to give a crude product. The crude product was dissolved in anhydrous 1,4 dioxane (5 mL), and then (3*S*)-3-aminotetrahydrofuran-2-one (1.20 eq, HBr salt) and Et_3_N (1.00 eq) were added. The resulting reaction mixture was degassed by purging with N_2_ 3 times, then heated to 100°C for 12 h under an N_2_ atmosphere. The reaction mixture was cooled to room temperature, diluted with H_2_O (10 mL) and extracted with EtOAc (3 x 10 mL). The combined organic layers were washed with brine (15 mL), dried over Na_2_SO_4_, filtered, and concentrated under reduced pressure to give a residue. The residue was purified by silica gel column chromatography to give the desired 3-oxo homoserine lactone as a white solid.

##### Procedure B

To a stirring solution of 2,2-dimethyl-1,3-dioxane-4,6-dione (Meldrum’s acid) (1.00 eq) and DMAP (1.05 eq) in DCM at 0°C was added DCC (1.10 eq) followed by the requisite carboxylic acid (1.00 eq). The resulting reaction mixture was allowed to warm to room temperature and stirred for 12 h. The reaction mixture was filtered through a pad of Celite to remove precipitated solids and concentrated under vacuum. The crude material was diluted with EtOAc (30 mL) and washed with cold 2N HCl (3 x 30 mL). The organic layer was separated, dried over Na_2_SO_4_, filtered and concentrated under reduced pressure to give a crude product. The crude product was dissolved in anhydrous 1,4 dioxane, and then (3*S*)-3-aminotetrahydrofuran-2-one (1.00 eq, HBr salt) and Et_3_N (1.00 eq) were added. The resulting reaction mixture was degassed by purging with N_2_ 3 times, then heated to 100°C for 12 h under an N_2_ atmosphere. The reaction mixture was cooled to room temperature, diluted with H_2_O (10 mL) and extracted with EtOAc (3 x 10 mL). The combined organic layers were washed with brine (15 mL), dried over Na_2_SO_4_, filtered, and concentrated under reduced pressure to give a residue. The residue was purified by silica gel column chromatography to give the desired 3-oxo homoserine lactone as a white solid.

#### (*S*)-3-oxo-*N*-(2-oxotetrahydrofuran-3-yl)hexanamide (BB0187)

Following Procedure A, elution with Petroleum ether/EtOAc = 1/1 afforded the title compound (285 mg, 29% yield, 97% purity by LC/MS) as a white solid. ^1^H NMR (400MHz, CDCl_3_) δ = 1H NMR (300 MHz, CDCl3) δ=7.85 (s, 1H), 4.67 (ddd, *J* = 6.7, 1H), 4.57 (td, *J* = 1.3, 4.2, 1H), 4.34 (m, 1H), 3.52 (s, 2H), 2.82 (m, 1H), 2.56 (t, *J* = 7.2, 2H) 2.33 (m, 1H), 1.64 (m, 2H), 0.98 (t, *J* = 6.5, 3H); MS (ESI) calculated for C_10_H_15_NO_4_: m/z = 213; found: m/z = 214 (M+H).

#### (*S*)-3-oxo-*N*-(2-oxotetrahydrofuran-3-yl)octanamide (BB0190)

Following Procedure A, elution with Petroleum ether/EtOAc = 1/1 afforded the title compound (220 mg, 37% yield, 99% purity by LC/MS) as a white solid. ^1^H NMR (400MHz, CDCl_3_) δ = 7.66 (s, br, 1H), 4.60 (m, 1H), 4.48 (t, *J* = 9.0 Hz, 1H), 4.29 (m, 1H), 3.47 (s, 2H), 2.77 (m, 1H), 2.53 (t, *J* = 7.3, 2H), 2.23, m, 1H), 1.61 (m, 2H), 1.31 (m, 4H), 0.90 (t, *J* = 6.6, 3H); MS (ESI) calculated for C_12_H_19_NO_4_: m/z = 241; found: m/z = 242 (M+H).

#### (*S*)-3-oxo-*N*-(2-oxotetrahydrofuran-3-yl)decanamide (BB0193)

Following Procedure A, gradient elution with Petroleum ether/EtOAc = 10/1 to 2/1 afforded the title compound (75 mg, 15% yield, 99% purity by LC/MS) as a white solid. ^1^H NMR (400MHz, CDCl_3_) δ = 7.68 (s, br, 1H), 4.60 (m, 1H), 4.48 (t, *J* = 9.1, 1H), 4.28 (m, 1H), 3.47 (s, 2H), 2.77 (m, 1H), 2.52 (t, *J* = 7.3, 2H), 2.22 (m, 1H), 1.59 (m, 2H), 1.27 (m, 8H), 0.88 (t, *J* = 6.2, 3H); MS (ESI) calculated for C_14_H_23_NO_4_: m/z = 269; found: m/z = 270 (M+H).

#### (*S*)-3-oxo-*N*-(2-oxotetrahydrofuran-3-yl)dodecanamide (BB0196)

Following Procedure A, gradient elution with Petroleum ether/EtOAc = 5/1 to 2/1 afforded the title compound (940 mg, 56% yield, 99% purity by LC/MS) as a white solid. ^1^H NMR (400MHz, CDCl_3_) δ = 7.67 (s, br, 1H), 4.58 (m, 1H), 4.48 (m, 1H), 4.28 (m, 1H), 3.47 (s, 2H), 2.75 (m, 1H), 2.52 (t, *J* = 7.3, 2H), 2.22 (m, 1H), 1.59 (m, 2H), 1.27 (m, 12H), 0.88 (t, *J* = 6.2, 3H); MS (ESI) calculated for C_16_H_27_NO_4_: m/z = 297; found: m/z = 298 (M+H).

#### (*S*)-3-oxo-*N*-(2-oxotetrahydrofuran-3-yl)tetradecanamide (BB0219)

Following Procedure B, gradient elution with Petroleum ether/EtOAc = 20/1 to 1/1 afforded the title compound (3.73 g, 46% yield, 98% purity by LC/MS) as a white solid. ^1^H NMR (400MHz, CDCl_3_) δ = 7.67 (d, J = 5.3, 1H), 4.60 (ddd, *J* = 6.8, 8.6, 11.4, 1H), 4.48 (dd, *J* = 9.2, 9.2, 1H), 4.28 (ddd, J = 5.9, 9.4, 11.0, 1H), 3.47 (s, 2H), 2.74 (m, 1H), 2.53 (t, *J* = 7.5, 2H), 2.23 (m, 1H), 1.59 (m, 2H), 1.26 (m, 16H), 0.91 (t, *J* = 6.8, 3H); MS (ESI) calculated for C_18_H_31_NO_4_: m/z = 325; found: m/z = 326 (M+H).

#### General procedure for the synthesis of 3-hydroxy homoserine lactones

To a stirring solution of 3-oxo homoserine lactone (1.00 eq) in DME (3 mL) at 0 °C was added NaBH_4_ (0.35 eq). The resulting reaction mixture was stirred at 0 °C for 2 h. The reaction mixture was concentrated under reduced pressure to give a residue. The residue was purified by silica gel column chromatography to give the desired 3-hydroxy homoserine lactone as a white solid.

#### 3-hydroxy-*N*-((*S*)-2-oxotetrahydrofuran-3-yl)hexanamide (BB0188)

Elution with EtOAc/CH_3_CN = 50/1 afforded the title compound (96 mg, 45% yield, 96% purity by LC/MS) as a white solid. ^1^H NMR (400MHz, CDCl_3_) δ = 6.64 (dd, br, *J* = 5.2, 20.0, 1H), 4.61 (m, 1H), 4.49 (t, *J* = 8.8, 2H), 4.30 (ddd, *J* = 6.0, 9.5, 11.0, 1H), 4.04 (br s, 1H), 3.18 (dd, *J* = 3.0, 17.1 Hz, 1H), 2.44 (m, 1H), 2.37 (m, 1H), 2.20(m, 1H), 1.53 - 1.39 (m, 4H), 0.94 (t, *J* = 6.8, 3H); MS (ESI) calculated for C_10_H_17_NO_4_: m/z = 215; found: m/z = 216 (M+H).

#### 3-hydroxy-*N*-((*S*)-2-oxotetrahydrofuran-3-yl)octanamide (BB0191)

Gradient elution with EtOAc/CH_3_CN = 80/1 to 70/1 afforded the title compound (96 mg, 32% yield, 99% purity by LC/MS) as a white solid. ^1^H NMR (400MHz, DMSO-d_6_) δ = 8.31 (s, br, 1H), 4.57 (m, 1H), 4.48 (m, 1H), 4.33 (m, 1H), 4.20 (m, 1H), 3.79 (m, 1H), 2.37 (m, 1H), - 2.16 (m, 3H), 1.40-1.11 (m, 8H), 0.86 (t, *J* = 6.6, 3H); MS (ESI) calculated for C_12_H_21_NO_4_: m/z = 243; found: m/z = 244 (M+H).

#### 3-hydroxy-*N*-((*S*)-2-oxotetrahydrofuran-3-yl)decanamide (BB0194)

Gradient elution with EtOAc/CH_3_CN = 80/1 to 70/1 afforded the title compound (106 mg, 38% yield, 99% purity by LC/MS) as a white solid. ^1^H NMR (400MHz, CDCl_3_) δ = 6.63 (d, br, *J* = 20.4, 1H), 4.59 (m, 1H), 4.49 (m, 1H), 4.30 (m, 1H), 4.03 (m, 1H), 3.15 (d, br, *J* = 14.0, 1H), 2.81 (m, 1H), 2.44 (m, 1H), 2.38 (m, 1H), 2.20 (m, 1H), 1.53 - 1.22 (m, 12H), 0.89 (t, *J* = 6.2, 3H); MS (ESI) calculated for C_14_H_25_NO_4_: m/z = 271; found: m/z = 272 (M+H).

#### 3-hydroxy-*N*-((*S*)-2-oxotetrahydrofuran-3-yl)dodecanamide (BB0197)

Elution with EtOAc/CH_3_CN = 100/1 afforded the title compound (137 mg, 47% yield, 99% purity by LC/MS) as a white solid. ^1^H NMR (400MHz, DMSO-d_6_) δ = 8.30 (dd, *J* = 8.1, 11.6, 1H), 4.54 (m, 2H), 4.33 (dt, *J* = 1.4, 8.8, 1H), 4.19 (m, 1H), 3.79 (m, 1H), 2.38 (m, 1H), 2.22-2.06 (m, 3H), 1.38-1.18 (m, 16H), 0.86 (t, *J* = 6.7, 3H); MS (ESI) calculated for C_16_H_29_NO_4_: m/z = 299; found: m/z = 300 (M+H).

### Statistical methods

In all experiments, values are the average of 3 biological replicates, each of which was assessed in 2 or 3 technical replicates as noted. For EC_50_ analyses, wildtype LasR was used as the control. EC_50_ values that appear in multiple experiments represent the mean from all experiments (Table 1, Figure 7, Figure S4 and Table S1). In these cases, 3 technical replicates of 3 biological replicates were assayed for every protein/molecule in each experiment. Biological replicates are defined as distinct samples analyzed on separate days. Technical replicates are defined as multiple measurements made on the same sample. Error bars represent standard deviations of the means. 2-tailed T-tests were performed to compare experimental groups. P values: *<.05, **<.01, ***<.0001

## Accession Number

The coordinates and structure factors will be deposited in the Protein Data Bank upon acceptance of the manuscript.

## Acknowledgements

We thank Dr. Fred Hughson and Dr. Philip Jeffrey for assistance with crystallography. We also thank Dr. Chari Smith and the entire Bassler group for insightful ideas about this research. This work was supported by the Howard Hughes Medical Institute, National Institutes of Health Grant 5R37GM065859, and National Science Foundation Grant MCB 1713731 (B.L.B.), NIGMS T32GM007388 (A.R.M), and a Jane Coffin Childs Memorial Fund for Biomedical Research Postdoctoral Fellowship (J.E.P.). The authors declare that they have no conflicts of interest with the contents of this article. The content is solely the responsibility of the authors and does not necessarily represent the official views of the National Institutes of Health.

## Author Contributions

A.R.M. performed the experiments and interpreted the data in Figures 1, 3, 5, 7, S2, S4 and Tables 1 and S1. J.E.P. performed the experiments and interpreted the data in Figures 2, 4, 6, 7, S1, and S3. B.R.H designed the synthesis for all the homoserine lactones used. B.L.B., A.R.M., and J.E.P conceived of the study. B.L.B coordinated the study and helped with interpretation of results. All of the authors contributed to the writing of the manuscript

## Supplementary Figures and Figure Legends

**Figure S1.**
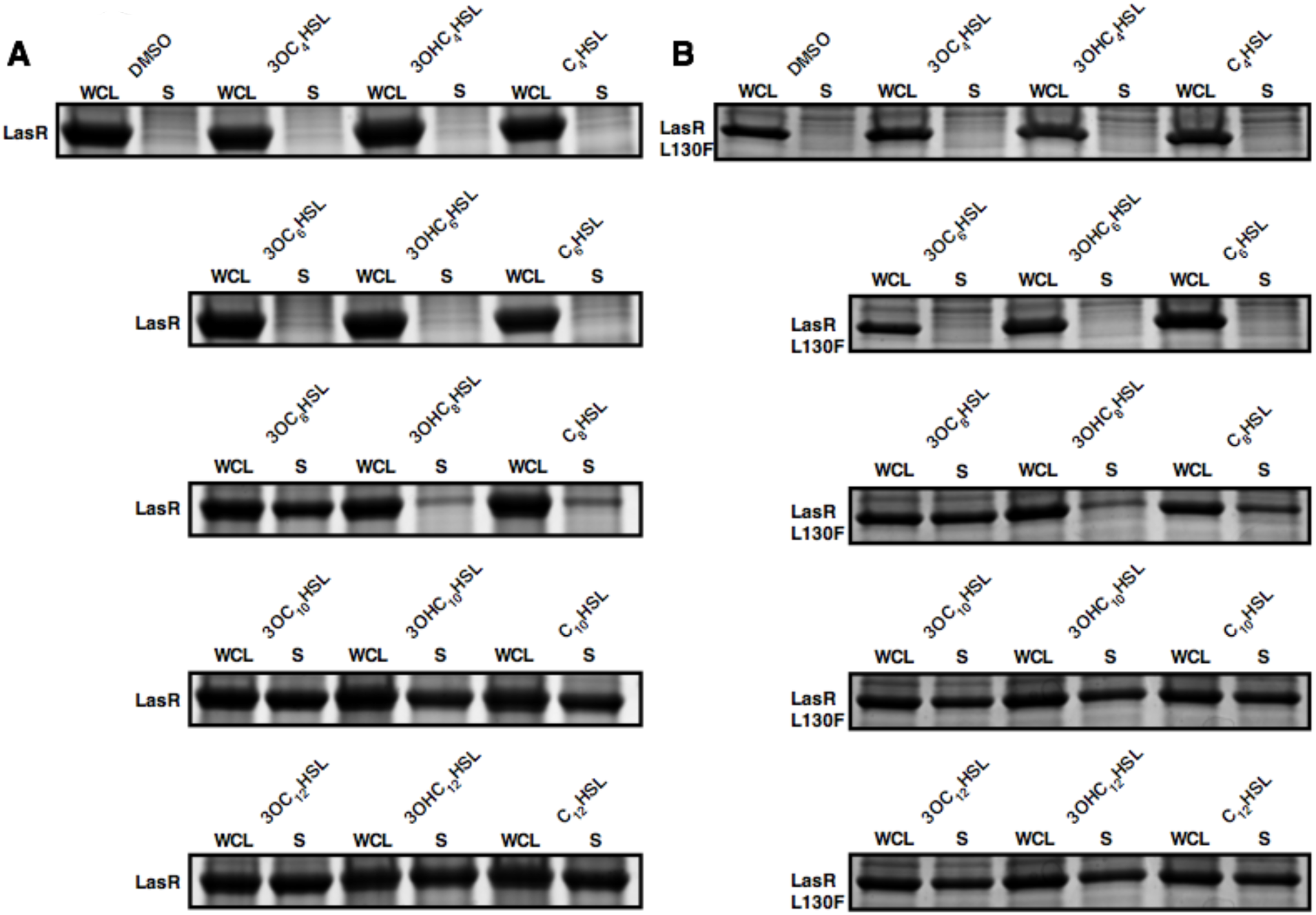
Long acyl chain HSLs solubilize the LasR LBD and the LasR LBD L130F. Comparison of wildtype LasR LBD protein levels in whole cell lysates (WCL) and in the soluble fractions (S) of *E. coli* cells harboring the DNA encoding the LasR LBD on a plasmid. B) As in panel A with LasR LBD L130F. In both panels, 1 mM IPTG was used for LasR induction and either 1% DMSO or 10 μM of the indicated HSL was added. See Figure 1, panel C of the main text for details.

**Figure S2.**
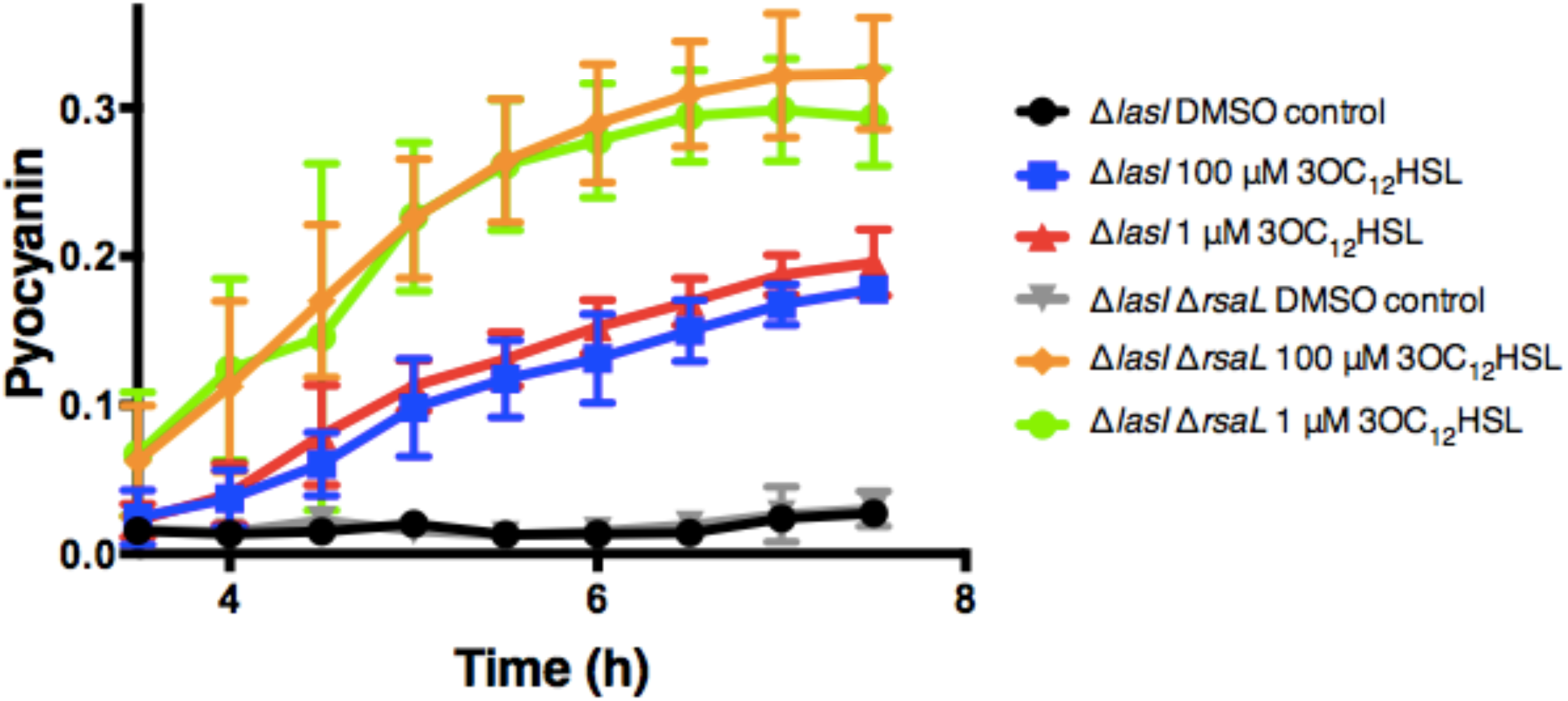
RsaL is required for inhibition of pyocyanin production by high HSL concentrations in P. aeruginosa. Pyocyanin production was measured spectophotometrically from *P. aeruginosa* Δ*lasI* and Δ*lasI* Δ*rsaL* strains over the growth curve following addition of DMSO, 1 μM 3OC_12_HSL, or 100 μM 3OC_12_HSL as indicated. Y-axis “Pyocyanin” is the amount of pyocyanin pigment (OD_695_ nm) over cell density (OD_600_ nm). Data show the mean of 3 biological replicates. Error bars denote standard deviations of the mean.

**Figure S3.**
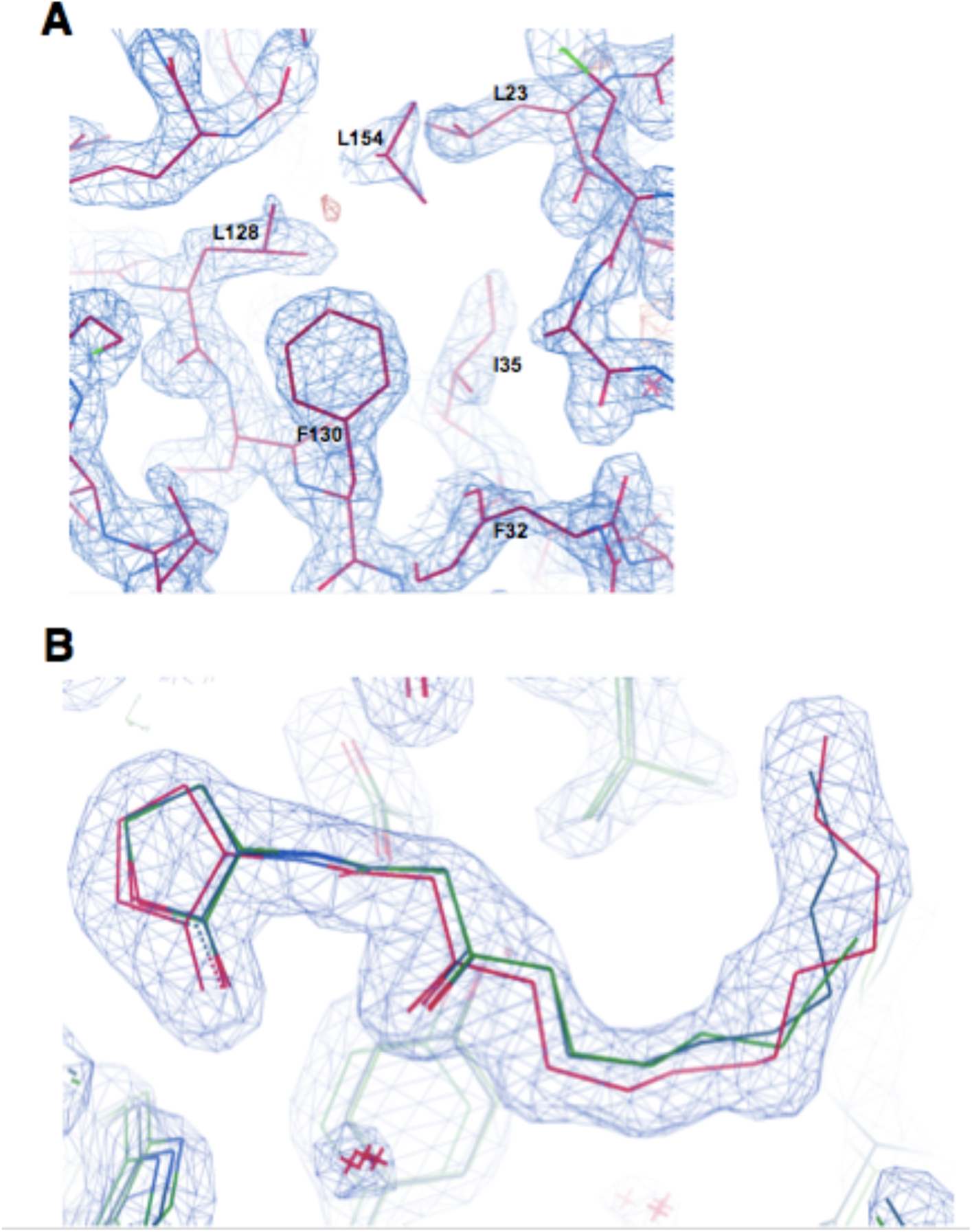
Electron density for LasR LBD L130F near residue F130 and the 3OC_14_HSL ligand. A) A simulated annealing omit map, contoured at 1σ, shows the electron density around LasR residue F130 and the surrounding hydrophobic residues. F130 interacts with L23, L30, F32, I35, L114, L118, L128, L151, and L154 in the LasR LBD L130F:3OC_14_HSL structure. The panel depicts the perspective highlighting the F130 interactions with L23, F32, I35, L128, and L154, and these residues are labeled. B) A simulated annealing omit map, contoured at 1σ, shows the electron density around the HSLs in the LasR LBD bound to 3OC_10_HSL (green), 3OC_14_HSL (red), or 3OC_12_HSL (blue, from data in Bottomley *et al*., 2007).

**Figure S4.**
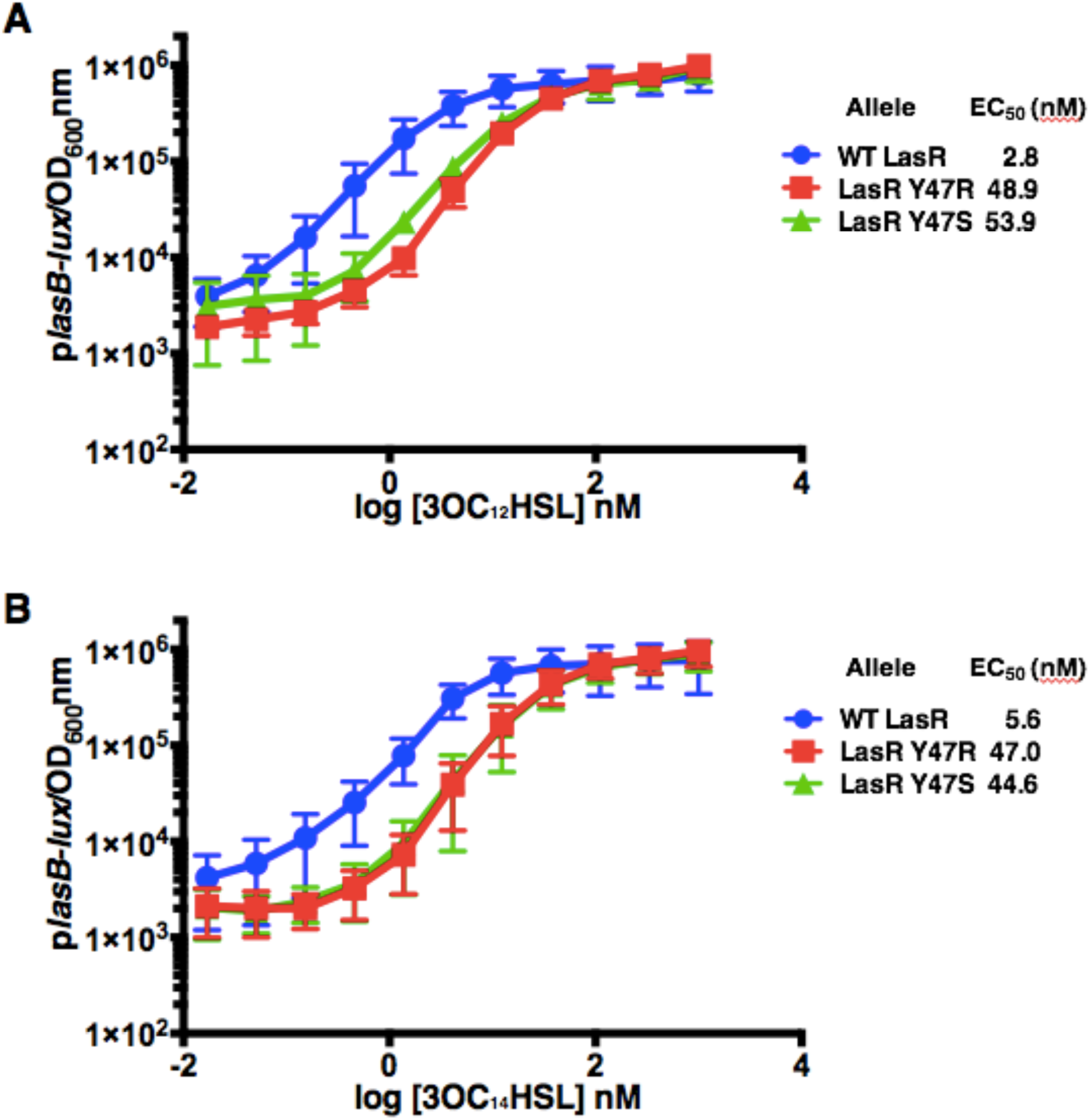
LasR Y47R and LasR Y47S have lower affinities for 3OC_12_HSL and 3OC_14_HSL than wildtype LasR. Bioluminescence from the p*lasB-lux* reporter driven by wildtype LasR (blue), LasR Y47R (red), and LasR Y47S (green) (See Figure 1, panel A of the main text for details). 3OC_12_HSL (A) or 3OC_14_HSL (B) were added at the designated concentrations. Two technical replicates were performed for each biological sample and 3 biological replicates were assessed. Error bars depict standard deviations of the mean.

**Table S1.**
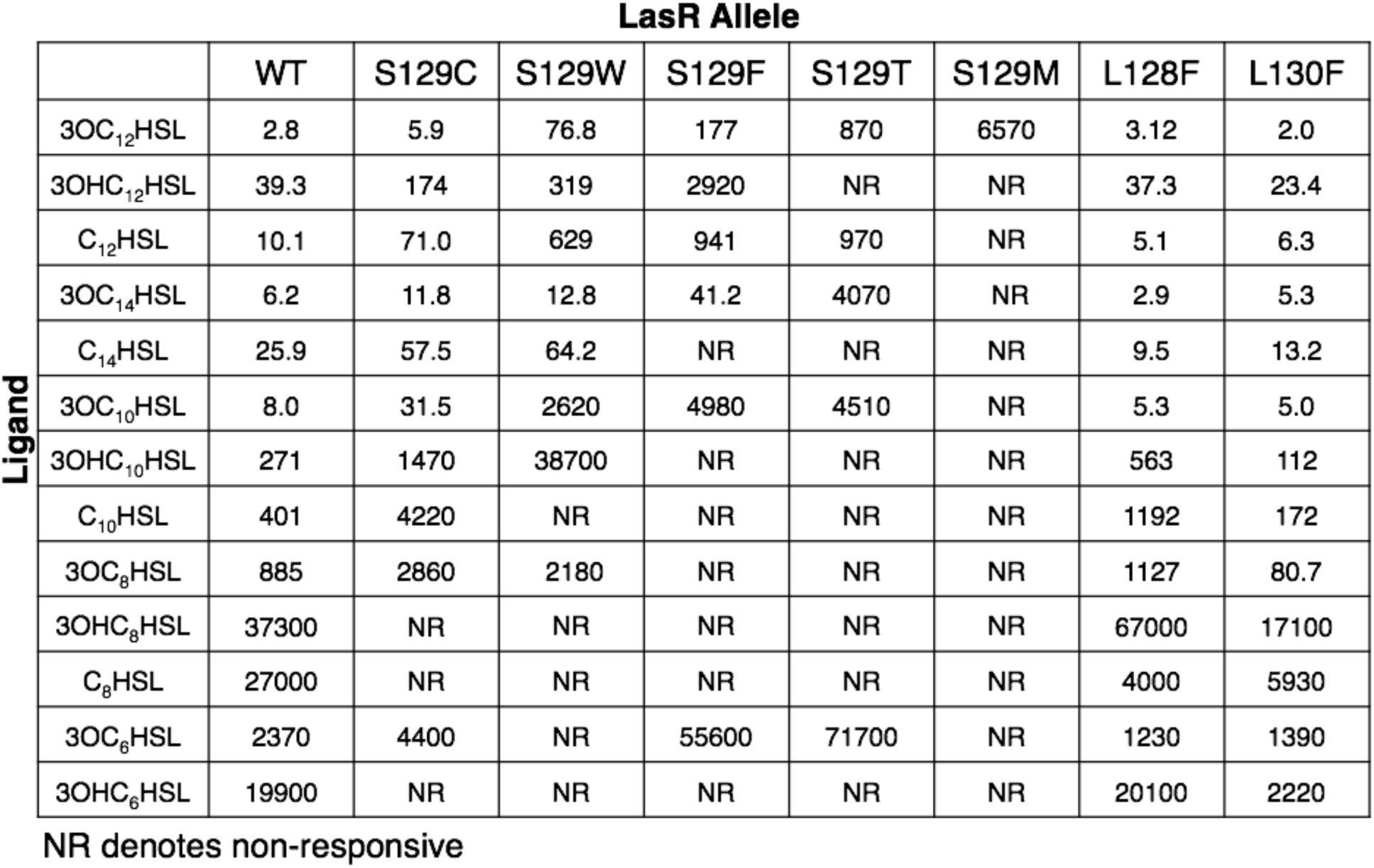
EC_50_ (nM) values for LasRand HSL compounds in the *plasB-lux* assay

| LasR Allele |  |  |  |  |  |  |  |  |  |
| --- | --- | --- | --- | --- | --- | --- | --- | --- | --- |
| Ligand |  | WT | S129C | S129W | S129F | S129T | S129M | L128F | L130F |
|  | 3OC<sub>12</sub>HSL | 2.8 | 5.9 | 76.8 | 177 | 870 | 6570 | 3.12 | 2.0 |
|  | 3OHC<sub>12</sub>HSL | 39.3 | 174 | 319 | 2920 | NR | NR | 37.3 | 23.4 |
|  | C<sub>12</sub>HSL | 10.1 | 71.0 | 629 | 941 | 970 | NR | 5.1 | 6.3 |
|  | 3OC<sub>14</sub>HSL | 6.2 | 11.8 | 12.8 | 41.2 | 4070 | NR | 2.9 | 5.3 |
|  | C<sub>14</sub>HSL | 25.9 | 57.5 | 64.2 | NR | NR | NR | 9.5 | 13.2 |
|  | 3OC<sub>10</sub>HSL | 8.0 | 31.5 | 2620 | 4980 | 4510 | NR | 5.3 | 5.0 |
|  | 3OHC<sub>10</sub>HSL | 271 | 1470 | 38700 | NR | NR | NR | 563 | 112 |
|  | C<sub>10</sub>HSL | 401 | 4220 | NR | NR | NR | NR | 1192 | 172 |
|  | 3OC<sub>8</sub>HSL | 885 | 2860 | 2180 | NR | NR | NR | 1127 | 80.7 |
|  | 3OHC<sub>8</sub>HSL | 37300 | NR | NR | NR | NR | NR | 67000 | 17100 |
|  | C<sub>8</sub>HSL | 27000 | NR | NR | NR | NR | NR | 4000 | 5930 |
|  | 3OC<sub>6</sub>HSL | 2370 | 4400 | NR | 55600 | 71700 | NR | 1230 | 1390 |
|  | 3OHC<sub>6</sub>HSL | 19900 | NR | NR | NR | NR | NR | 20100 | 2220 |

NR denotes non-responsive

## Literature Cited

Adams, P. D., Afonine, P. V., Bunkóczi, G., Chen, V. B., Echols, N., Headd, J. J., … Zwart, P. H. (2011). The Phenix software for automated determination of macromolecular structures. Methods, 55(1), 94–106. doi:10.1016/j.ymeth.2011.07.005

Afonine, P. V., Grosse-Kunstleve, R. W., Echols, N., Headd, J. J., Moriarty, N. W., Mustyakimov, M., … Adams, P. D. (2012). Towards automated crystallographic structure refinement with phenix.refine. Acta Crystallogr D Biol Crystallogr, 68(Pt 4), 352–367. doi:10.1107/S0907444912001308

Albus, A. M., Pesci, E. C., RunyenJanecky, L. J., West, S. E. H., & Iglewski, B. H. (1997). Vfr controls quorum sensing in Pseudomonas aeruginosa. Journal of Bacteriology, 179(12), 3928–3935.

Baron, S. S., & Rowe, J. J. (1981). Antibiotic action of pyocyanin. Antimicrobial Agents and Chemotherapy, 20(6), 814–820. doi:10.1128/aac.20.6.814

Bassler, B. L., Wright, M., Showalter, R. E., & Silverman, M. R. (1993). Intercellular Signaling in Vibrio-Harveyi - Sequence and Function of Genes Regulating Expression of Luminescence. Molecular Microbiology, 9(4), 773–786. doi:DOI 10.1111/j.1365-2958.1993.tb01737.x

Borlee, B. R., Geske, G. D., Blackwell, H. E., & Handelsman, J. (2010). Identification of Synthetic Inducers and Inhibitors of the Quorum-Sensing Regulator LasR in Pseudomonas aeruginosa by High-Throughput Screening. Applied and Environmental Microbiology, 76(24), 8255–8258. doi:10.1128/aem.00499-10

Bottomley, M. J., Muraglia, E., Bazzo, R., & Carfi, A. (2007). Molecular insights into quorum sensing in the human pathogen Pseudomonas aeruginosa from the structure of the virulence regulator LasR bound to its autoinducer. Journal of Biological Chemistry, 282(18), 13592–13600. doi:10.1074/jbc.M700556200

Brint, J. M., & Ohman, D. E. (1995). Sythnesis of multiple exoproducts in Pseudomonas-aeruginosa is under the control of the RhlR-RHlI, another set of regulators in strain PAO1 with homology to the autoinducer-responsive LuxR-LuxI Family Journal of Bacteriology, 177(24), 7155–7163. doi:10.1128/jb.177.24.7155-7163.1995

Cabeen, M. T. (2014). Stationary Phase-Specific Virulence Factor Overproduction by a IasR Mutant of Pseudomonas aeruginosa. Plos One, 9(2), 9. doi:10.1371/journal.pone.0088743

Cao, J. G., & Meighen, E. A. (1989). Purification and structural identification of an autoinducer for the luminescence system of Vibrio harveyi. J Biol Chem, 264(36), 21670–21676.

Chen, G., Swem, L. R., Swem, D. L., Stauff, D. L., O’Loughlin, C. T., Jeffrey, P. D., … Hughson, F. M. (2011). A strategy for antagonizing quorum sensing. Mol Cell, 42(2), 199–209. doi:10.1016/j.molcel.2011.04.003

Chhabra, S. R., Harty, C., Hooi, D. S. W., Daykin, M., Williams, P., Telford, G., … Bycroft, B. W. (2003). Synthetic analogues of the bacterial signal (quorum sensing) molecule N-(3-oxododecanoyl)-L-homoserine lactone as immune modulators. Journal of Medicinal Chemistry, 46(1), 97–104. doi:10.1021/jm020909n

Churchill, M. E. A., & Chen, L. L. (2011). Structural Basis of Acyl-homoserine Lactone-Dependent Signaling. Chemical Reviews, 111(1), 68–85. doi:10.1021/cr1000817

DeLano, W. L. (2009). PyMOL molecular viewer: Updates and refinements. Abstracts of Papers of the American Chemical Society, 238, 1.

Eberhard, A., Burlingame, A. L., Eberhard, C., Kenyon, G. L., Nealson, K. H., & Oppenheimer, N. J. (1981). Structural identification of autoinducer of Photobacterium fischeri luciferase. Biochemistry, 20(9), 2444–2449. doi:10.1021/bi00512a013

Emsley, P., & Cowtan, K. (2004). Coot: model-building tools for molecular graphics. Acta Crystallogr D Biol Crystallogr, 60(Pt 12 Pt 1), 2126–2132. doi:10.1107/S0907444904019158

Emsley, P., Lohkamp, B., Scott, W. G., & Cowtan, K. (2010). Features and development of Coot. Acta Crystallogr D Biol Crystallogr, 66(Pt 4), 486–501. doi:10.1107/S0907444910007493

Engebrecht, J., Nealson, K., & Silverman, M. (1983). Bacterial bioluminescence: isolation and genetic analysis of functions from Vibrio fischeri. Cell, 32(3), 773–781. doi:10.1016/0092-8674(83)90063-6

Feltner, J. B., Wolter, D. J., Pope, C. E., Groleau, M. C., Smalley, N. E., Greenberg, E. P., … Dandekar, A. A. (2016). LasR Variant Cystic Fibrosis Isolates Reveal an Adaptable Quorum-Sensing Hierarchy in Pseudomonas aeruginosa. Mbio, 7(5), 9. doi:10.1128/mBio.01513-16

Freeman, J. A., Lilley, B. N., & Bassler, B. L. (2000). A genetic analysis of the functions of LuxN: a two-component hybrid sensor kinase that regulates quorum sensing in Vibrio harveyi. Molecular Microbiology, 35(1), 139–149. doi:10.1046/j.1365-2958.2000.01684.x

Gambello, M. J., & Iglewski, B. H. (1991). Cloning and characterization of the Pseudomonas aeruginosa lasR gene, a transcriptional activator of elastase expression. J Bacteriol, 173(9), 3000–3009.

Gao, J., Ma, A. Z., Zhuang, X. L., & Zhuang, G. Q. (2014). An N-Acyl Homoserine Lactone Synthase in the Ammonia-Oxidizing Bacterium Nitrosospira multiformis. Applied and Environmental Microbiology, 80(3), 951–958. doi:10.1128/aem.03361-13

Gerdt, J. P., McInnis, C. E., Schell, T. L., & Blackwell, H. E. (2015). Unraveling the contributions of hydrogen-bonding interactions to the activity of native and non-native ligands in the quorum-sensing receptor LasR. Organic & Biomolecular Chemistry, 13(5), 1453–1462. doi:10.1039/c4ob02252a

Gerdt, J. P., McInnis, C. E., Schell, T. L., Rossi, F. M., & Blackwell, H. E. (2014). Mutational Analysis of the Quorum-Sensing Receptor LasR Reveals Interactions that Govern Activation and Inhibition by Nonlactone Ligands. Chemistry & Biology, 21(10), 1361–1369.

Gerdt, J. P., Wittenwyler, D. M., Combs, J. B., Boursier, M. E., Brummond, J. W., Xu, H., & Blackwell, H. E. (2017). Chemical Interrogation of LuxR-type Quorum Sensing Receptors Reveals New Insights into Receptor Selectivity and the Potential for Interspecies Bacterial Signaling. Acs Chemical Biology, 12(9), 2457–2464. doi:10.1021/acschembio.7b00458

Hanzelka, B. L., Parsek, M. R., Val, D. L., Dunlap, P. V., Cronan, J. E., & Greenberg, E. P. (1999). Acylhomoserine lactone synthase activity of the Vibrio fischeri AinS protein. Journal of Bacteriology, 181(18), 5766–5770.

Hawver, L. A., Jung, S. A., & Ng, W. L. (2016). Specificity and complexity in bacterial quorum-sensing systems. FEMS Microbiol Rev, 40(5), 738–752. doi:10.1093/femsre/fuw014

Ke, X. B., Miller, L. C., & Bassler, B. L. (2015). Determinants governing ligand specificity of the Vibrio harveyi LuxN quorum-sensing receptor. Molecular Microbiology, 95(1), 127–142. doi:10.1111/mmi.12852

Kukavica-Ibrulj, I., Bragonzi, A., Paroni, M., Winstanley, C., Sanschagrin, F., O’Toole, G. A., & Levesque, R. C. (2008). In vivo growth of Pseudomonas aeruginosa strains PAO1 and PA14 and the hypervirulent strain LESB58 in a rat model of chronic lung infection. Journal of Bacteriology, 190(8), 2804–2813. doi:10.1128/jb.01572-07

Latifi, A., Winson, M. K., Foglino, M., Bycroft, B. W., Stewart, G. S. A. B., Lazdunski, A., & Williams, P. (1995). Multiple Homologs of Luxr and Luxl Control Expression of Virulence Determinants and Secondary Metabolites through Quorum Sensing in Pseudomonas-Aeruginosa Pao1. Molecular Microbiology, 17(2), 333–343. doi:DOI 10.1111/j.1365-2958.1995.mmi_17020333.x

Mellbye, B. L., Spieck, E., Bottomley, P. J., & Sayavedra-Soto, L. A. (2017). Acyl-Homoserine Lactone Production in Nitrifying Bacteria of the Genera Nitrosospira, Nitrobacter, and Nitrospira Identified via a Survey of Putative Quorum-Sensing Genes. Applied and Environmental Microbiology, 83(22), 13. doi:10.1128/aem.01540-17

Michael, B., Smith, J. N., Swift, S., Heffron, F., & Ahmer, B. M. (2001). SdiA of Salmonella enterica is a LuxR homolog that detects mixed microbial communities. J Bacteriol, 183(19), 5733–5742. doi:10.1128/JB.183.19.5733-5742.2001

Minor, W., Cymborowski, M., Otwinowski, Z., & Chruszcz, M. (2006). HKL-3000: the integration of data reduction and structure solution--from diffraction images to an initial model in minutes. Acta Crystallogr D Biol Crystallogr, 62(Pt 8), 859–866. doi:10.1107/S0907444906019949

Mukherjee, S., Moustafa, D., Smith, C. D., Goldberg, J. B., & Bassler, B. L. (2017). The RhlR quorum-sensing receptor controls Pseudomonas aeruginosa pathogenesis and biofilm development independently of its canonical homoserine lactone autoinducer. Plos Pathogens, 13(7), 25. doi:10.1371/journal.ppat.1006504

Nasser, W., & Reverchon, S. (2007). New insights into the regulatory mechanisms of the LuxR family of quorum sensing regulators. Analytical and Bioanalytical Chemistry, 387(2), 381–390. doi:10.1007/s00216-006-0702-0

Nguyen, Y., Nguyen, N. X., Rogers, J. L., Liao, J., MacMillan, J. B., Jiang, Y., & Sperandio, V. (2015). Structural and mechanistic roles of novel chemical ligands on the SdiA quorum-sensing transcription regulator. MBio, 6(2). doi:10.1128/mBio.02429-14

O’Loughlin, C. T., Miller, L. C., Siryaporn, A., Drescher, K., Semmelhack, M. F., & Bassler, B. L. (2013). A quorum-sensing inhibitor blocks Pseudomonas aeruginosa virulence and biofilm formation. Proceedings of the National Academy of Sciences of the United States of America, 110(44), 17981–17986. doi:10.1073/pnas.1316981110

Paczkowski, J. E., Mukherjee, S., McCready, A. R., Cong, J. P., Aquino, C. J., Kim, H., … Bassler, B. L. (2017). Flavonoids Suppress Pseudomonas aeruginosa Virulence through Allosteric Inhibition of Quorum-sensing Receptors. J Biol Chem, 292(10), 4064–4076. doi:10.1074/jbc.M116.770552

Papenfort, K., & Bassler, B. L. (2016). Quorum sensing signal-response systems in Gram-negative bacteria. Nature Reviews Microbiology, 14(9), 576–588. doi:10.1038/nrmicro.2016.89

Pearson, J. P., Gray, K. M., Passador, L., Tucker, K. D., Eberhard, A., Iglewski, B. H., & Greenberg, E. P. (1994). Structure of the autoinducer required for expression of Pseudomonas aeruginosa virulence genes. Proc Natl Acad Sci U S A, 91(1), 197–201. doi:10.1073/pnas.91.1.197

Pearson, J. P., Pesci, E. C., & Iglewski, B. H. (1997). Roles of Pseudomonas aeruginosa las and rhl quorum-sensing systems in control of elastase and rhamnolipid biosynthesis genes. Journal of Bacteriology, 179(18), 5756–5767.

Rampioni, G., Schuster, M., Greenberg, E. P., Bertani, I., Grasso, M., Venturi, V., … Leoni, L. (2007). RsaL provides quorum sensing homeostasis and functions as a global regulator of gene expression in Pseudomonas aeruginosa. Molecular Microbiology, 66(6), 1557–1565. doi:10.1111/j.1365-2958.2007.06029.x

Schuster, M., & Greenberg, E. P. (2007). Early activation of quorum sensing in Pseudomonas aeruginosa reveals the architecture of a complex regulon. Bmc Genomics, 8, 11. doi:10.1186/1471-2164-8-287

Schuster, M., Urbanowski, M. L., & Greenberg, E. P. (2004). Promoter specificity in Pseudomonas aeruginosa quorum sensing revealed by DNA binding of purified LasR. Proceedings of the National Academy of Sciences of the United States of America, 101(45), 15833–15839. doi:10.1073/pnas.0407229101

Schweizer, H. P. (1991). Escherichia-Pseudomonas shuttle vectors derived from pUC18/19. Gene, 97(1), 109–121. doi:10.1016/0378-1119(91)90016-5

Simon, R., Priefer, U., & Puhler, A. (1983). A Broad Host Range Mobilization System for Invivo Genetic-Engineering - Transposon Mutagenesis in Gram-Negative Bacteria. Bio-Technology, 1(9), 784–791. doi:DOI 10.1038/nbt1183-784

Sitnikov, D. M., Schineller, J. B., & Baldwin, T. O. (1996). Control of cell division in Escherichia coli: regulation of transcription of ftsQA involves both rpoS and SdiA-mediated autoinduction. Proc Natl Acad Sci U S A, 93(1), 336–341.

Smalley, N. E., An, D. D., Parsek, M. R., Chandler, J. R., & Dandekar, A. A. (2015). Quorum Sensing Protects Pseudomonas aeruginosa against Cheating by Other Species in a Laboratory Coculture Model. Journal of Bacteriology, 197(19), 3154–3159. doi:10.1128/jb.00482-15

Tashiro, Y., Yawata, Y., Toyofuku, M., Uchiyama, H., & Nomura, N. (2013). Interspecies Interaction between Pseudomonas aeruginosa and Other Microorganisms. Microbes and Environments, 28(1), 13–24. doi:10.1264/jsme2.ME12167

Vannini, A., Volpari, C., Gargioli, C., Muraglia, E., Cortese, R., De Francesco, R., … Di Marco, S. (2002). The crystal structure of the quorum sensing protein TraR bound to its autoinducer and target DNA. Embo Journal, 21(17), 4393–4401. doi:10.1093/emboj/cdf459

You, Y. S., Marella, H., Zentella, R., Zhou, Y. Y., Ulmasov, T., Ho, T. H. D., & Quatrano, R. S. (2006). Use of bacterial quorum-sensing components to regulate gene expression in plants. Plant Physiology, 140(4), 1205–1212. doi:10.1104/pp.105.074666

Zhang, R. G., Pappas, K. M., Pappas, T., Brace, J. L., Miller, P. C., Oulmassov, T., … Joachimiak, A. (2002). Structure of a bacterial quorum-sensing transcription factor complexed with pheromone and DNA. Nature, 417(6892), 971–974. doi:10.1038/nature00833

Zhu, J., & Winans, S. C. (2001). The quorum-sensing transcriptional regulator TraR requires its cognate signaling ligand for protein folding, protease resistance, and dimerization. Proc Natl Acad Sci U S A, 98(4), 1507–1512. doi:10.1073/pnas.98.4.1507

